# Single-Nucleotide m^6^A Mapping Uncovers Redundant YTHDF Function in Planarian Progenitor Fate Selection

**DOI:** 10.1101/2025.03.03.641144

**Authors:** Yarden Yesharim, Ophir Shwarzbard, Jenny Barboy-Smoliarenko, Ran Shachar, Amrutha Palavalli, Hanh Thi-Kim Vu, Schraga Schwartz, Omri Wurtzel

## Abstract

Cell fate decisions require tight regulation of gene expression. In planarians—highly regenerative flatworms—the mRNA modification N^6^-methyladenosine (m^6^A) modulates progenitor production and fate. However, the mechanisms governing m^6^A deposition in the planarian transcriptome—and the role of their expanded family of YTHDF m^6^A reader proteins in orchestrating biological functions—remain unclear. Here, we generated the first single-nucleotide resolution map of m^6^A in planarians, and revealed that simple sequence rules guide m^6^A deposition, facilitating the flexible evolutionary gain and loss of these marks. Functional analyses of the five planarian m^6^A readers revealed that while individual reader expression is dispensable, together, the planarian readers regulate the production of specific progenitor lineages and overall body size. Collectively, our findings uncover a robust, redundant regulatory architecture, characterized by multiple m^6^A sites per gene and coordinated m^6^A reader expression. This architecture is essential for proper lineage resolution and provides insights into the evolutionary dynamics of the m^6^A landscape.

## Introduction

Regeneration is a highly dynamic process that demands coordination of gene expression programs for new cell production and recovery of damaged tissues^1–3^. In recent years, post-transcriptional modifications have emerged as important regulators of such dynamic gene expression changes^4,5^. Among these modifications, N^6^-methyladenosine (m^6^A) is the most prevalent across the transcriptome^6,7^, and has critical regulatory roles in diverse developmental systems^7–15^. In planarians – highly regenerative flatworms – m^6^A is essential for the production of progenitor cells in specific lineages, such as the intestine^4^, and for suppressing the excessive production of neural-like progenitor cells^4^, echoing observations in other organisms where m^6^A modulates key developmental processes^13,16–19^.

Deciphering how m^6^A influences complex developmental programs is highly challenging for several reasons^20^. First, m^6^A is pervasive. In planarians, approximately 7,000 different transcripts are modified, displaying variable levels of m^6^A stoichiometries, and expressed across different cell types and states^4^. Second, m^6^A functions are mediated by a diverse set of reader proteins, the largest being the YTH-family proteins (YTHDF)^7,8,10^. Notably, planarians possess an expanded family of m^6^A readers – at least five YTHDF proteins – complicating efforts to determine their individual and combined roles^4^. Third, conventional approaches for functional analysis, such as inhibiting genes encoding core components of the methyltransferase complex (MTC) or the gene encoding the nuclear m^6^A reader YTHDC-1, result in severe phenotypes including lethality and complete loss of regenerative ability, thereby masking the roles of other m^6^A readers^4,21^.

The limited understanding of the rules governing m^6^A deposition further complicates the picture. Two major models have been proposed in vertebrate systems: one in which m^6^A is deposited pervasively on compatible sequence motifs but is excluded from regions that are inaccessible to the MTC (for example, in vertebrates regions bound by the exon junction complex; EJC)^22–25^, and another, where deposition is more finely regulated based on organismal requirements^5,26,27^. In planarians, our recent study has suggested that the sequence enriched at m^6^A sites is remarkably simple (GAC motif)^4^ compared to the more complex DRACH motif found in other organisms (e.g., humans)^6^. Whether an exclusion mechanism similar to that observed in vertebrates^22^ operates in planarians—and how this might influence the evolutionary dynamics of m^6^A site gain and loss—remains an open question. The availability of single-base resolution profiling methods based on sequencing, such as GLORI, facilitates a finer analysis of m^6^A sequence preference and their conservation^28,29^.

In addition to the challenges of understanding m^6^A installation^28^, the interpretation of this mark by the m^6^A readers is critical for understanding the regulation of its targets^7,12^. While vertebrates have three YTHDF paralogs^10^ and *Drosophila* has only one^11^, planarian genomes have at least five *ythdf*s genes^4^. Whether these factors function in a redundant manner, as suggested by several studies in vertebrates^9,12,14^, or exert specialized, state or lineage-specific regulatory roles^30^, is unknown. Work in other systems provided evidence supporting both models: some studies have demonstrated that individual YTHDF proteins target distinct sets of m^6^A-modified transcripts^31–35^, while others have found extensive overlap in their functions^9,12,14^. Resolving this ambiguity in planarians is essential for understanding how m^6^A modifications are translated into specific cellular outcomes during new cell production.

To address these challenges, we mapped m^6^A sites in planarians at single-nucleotide resolution, identified 19,328 m^6^A sites across the planarian transcriptome, and characterized the underlying sequence rules governing their deposition. Notably, m^6^A sites appear to be installed independently on each gene, suggesting that genes can gain or lose m^6^A modifications without compromising the functionality of existing sites. Our analysis supports a model in which the evolutionary gain and loss of m^6^A sites is subject to minimal sequence constraints, thereby allowing flexible m^6^A pattern formation across transcripts. Furthermore, by examining the expression patterns and functional contributions of the expanded family of planarian YTHDF proteins, we found that their expression largely overlaps. Moreover, only the simultaneous suppression of multiple YTHDF-encoding genes produced striking phenotypes – animals displaying reduced body size and having a diminished pool of parenchymal progenitors and *cathepsin*^+^ cell types. These findings suggested that planarian YTHDFs act, to a large extent, redundantly to promote the production of specific lineages, thereby ensuring proper tissue homeostasis. Collectively, our work advances the understanding of the mechanisms governing m^6^A deposition and recognition, and also provides new insights into the evolutionary dynamics of m^6^A regulation in this regenerative organism and raises new mechanistic questions regarding m^6^A deposition and readout.

## Results

### Single-nucleotide mapping of planarian m^6^A sites using GLORI

m^6^A is abundant across the planarian transcriptome^4^, yet previous profiling efforts lacked the resolution required to elucidate the principles governing its deposition. To address this gap, we used GLORI^28,29^, a method that identifies m^6^A sites at a single-nucleotide resolution. RNA was analyzed from control samples, having normal m^6^A levels, and *kiaa1429* (RNAi) animals, in which m^6^A levels are reduced due to inhibited methyltransferase complex (MTC) activity^4^. GLORI selectively converts adenosines to inosines, but not m^6^A^28^, enabling quantification of the m^6^A-to-A ratio across the transcriptome. Following GLORI conversion, we prepared and sequenced cDNA libraries, and mapped them to the planarian genome^36^ (Methods). For each adenosine in the transcriptome, we calculated an m^6^A score by dividing the number of reads having A mapped to the position by the number of reads having either A or G (the sequencing product of inosine) mapped to the position (Fig 1A-B; Table S1-2).

**Figure 1.**
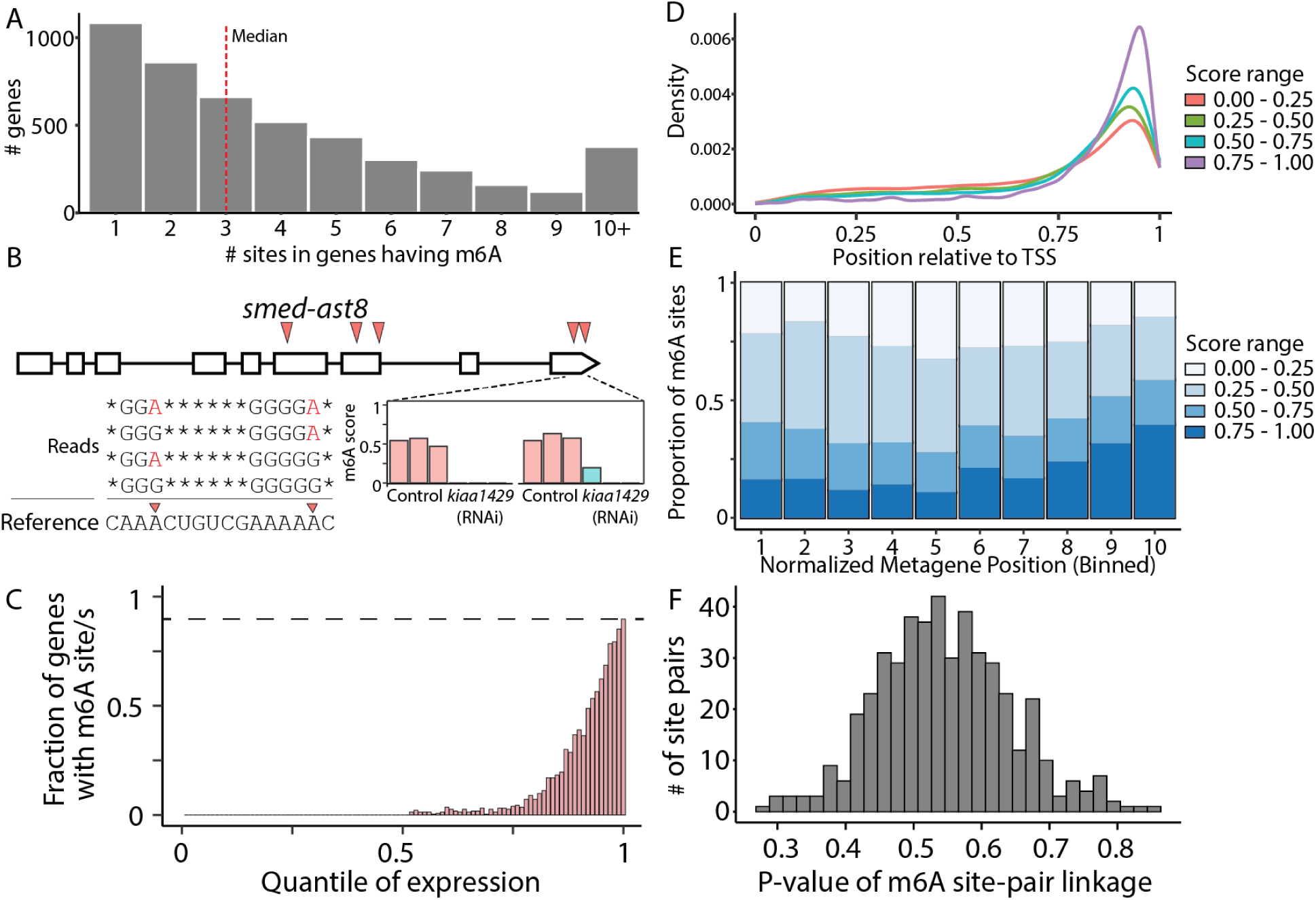
Characterization of planarian m6A distribution using GLORI. (A) The number of detected m^6^A sites (Methods) per gene is shown. The single nucleotide resolution m^6^A mapping facilitated detection of multiple sites per gene. (B) Mapping of m^6^A sites across the gene *ast8*, identified five m^6^A sites (top; red arrowheads), including two sites that were closely adjacent (12 nt). Sequencing reads spanning both the adjacent sites (bottom-left) show how m^6^A sites were detected: Reads containing adenosines (red letters) indicate that a methyl group protected the nucleotide from the GLORI treatment. For each detected site, a score was calculated (bottom-right). Shown are the scores in three biological replicates in control animals (red bars) and in animals depleted of m^6^A (*kiaa1429* (RNAi)), due to suppression of the MTC ^4^. (C) Shown is the fraction of genes having m^6^A site as a function of the expression percentile (Methods). (D) Meta-gene analysis of m^6^A localization showed a strong 3’-end bias of m^6^A sites, regardless of the m^6^A score. (E) Distribution of m^6^A scores at different regions across the length of the transcript (bins 1 to 10 indicate relative positions from the 5’ to 3’ ends, respectively). Sites near the 3’ end were modified more frequently compared to other m^6^A sites across the transcript. (F) Distribution of p-values calculated for assessing linkage in m^6^A installation at nearby sites. The distribution of p-values supported the interpretation that there was no association in installation of m^6^A in nearby sites, and that m^6^A was installed independently at every site.

We identified 19,328 m^6^A sites across 4,718 genes (Fig 1A-B), with conversion rates exceeding 10% in any sample (Fig S1A; Table S1; Control and *kiaa1429* (RNAi), 15309 and 11274 m6A sites, respectively). In control samples, 5,264 sites (27.2%) displayed a median m^6^A score greater than 0.5, while only 1,818 sites (9.4%) met this criterion in the *kiaa1429* (RNAi) samples (Table S1; Fig S1B). This reduction in the number of detectable m^6^A sites demonstrated the impact of inhibiting this MTC component on m^6^A levels (Fig S1B). Further examination of m^6^A sites in genes expressed in all samples revealed a median m^6^A score of 0.88 and 0.45, in control and *kiaa1429* (RNAi) samples, respectively (Fig S1C-D; n = 2,284; Site read coverage > 10; Methods). Detection of m^6^A sites using GLORI required sufficient gene expression. To evaluate how expression levels influenced m^6^A site detection, we utilized sequencing libraries from the same RNA used for GLORI, which we left untreated. Normalized gene expression values were calculated from the untreated libraries (Methods) and assigned to expression percentiles (1-100; Fig S1E). Using the GLORI libraries, we analyzed the number of m^6^A sites detected within genes assigned to each expression percentile (Fig 1C, S1F). The detection of m^6^A sites was strongly correlated with gene expression levels. For instance, over 50% of the genes were not expressed at all, and therefore lacked detectable m^6^A sites (Fig S1E-F). Furthermore, only 2.7% of the genes in the 70th percentile had m^6^A sites detectable across biological replicates (Fig 1C, S1E; Table S1). Detection increased with expression level, with 90% of the genes in the top expression percentile containing detectable m^6^A sites (Fig 1C). This could either be a characteristic of the most highly expressed genes or, alternatively, suggest that m^6^A modifications are extremely abundant but remain undetectable in transcripts with insufficient expression levels during m^6^A profiling assays.

m^6^A sites were preferentially found near the 3’ end of transcripts (Fig 1D), a pattern that persisted in both multi-exon and single-exon genes (Fig 1D, S1G; Table S1). Moreover, m^6^A sites located towards the 3’ end had a higher score compared to other m^6^A sites (Fig 1E; Student’s t-test p = 7.87e-54; Methods). m^6^A sites were even detected in non-polyadenylated, single-exon histone transcripts, indicating that this preference was not necessarily dependent on polyadenylation associated factors, polyadenylation sequence signals, or the splicing machinery (Fig S1G-H; Table S1).

Single-nucleotide resolution detection of m^6^A revealed that individual genes often contained multiple, closely spaced m^6^A sites (Fig 1A-B). The median distance between the highest scoring m^6^A site in a gene and the nearest secondary site was 18 nucleotides (Fig S1I-J; Table S1). This m^6^A site proximity could be a consequence of the MTC activity on several nearby positions on the same molecule (i.e., single-molecule linkage, e.g., via coupled deposition), or the independent activity of the MTC on different molecules (i.e., regional linkage, e.g., guided by *cis* elements). We examined sequencing reads spanning two adjacent m^6^A sites, and determined whether the modification of one site influenced the likelihood of methylation at the other site (Fig 1B, F, S1K). We selected m^6^A site pairs for this analysis that were separated by up-to 40 nt, and exhibited methylation levels between 0.4 and 0.7 (n = 479). Our results indicated no evidence of linkage between such adjacent sites at the single molecule resolution (Fig 1F, S1K; Methods), suggesting that methylation at one site did not alter the probability of methylation at a neighboring site on the same molecule. These results indicated that methylation of adjacent sites were a consequence of independent events of MTC activity, and not an outcome of processive MTC activity at nearby sites. Interestingly, despite lack of linkage in methylation of adjacent sites, we found a moderate correlation (Pearson r = 0.49, p-value < 2.2e-16; Methods) between the m^6^A scores of adjacent sites, meaning that if one site had a high m^6^A score, a nearby site was more likely to have a high score as well (Fig S1L-M). Altogether, this suggested that local characteristics of the transcript contributed to the level of methylation of nearby sites, in line with recent discoveries^37^.

### Analysis of sequence preferences associated with m^6^A installation

We next examined the sequences surrounding m^6^A sites and found that even sequences deviating from the canonical installation motif (i.e., DRACH^6^) can serve as excellent MTC targets, provided they conform to specific rules (Fig 2A-D). Essentially all m^6^A sites were followed by either a cytosine or uracil, with strong depletion of A and G (Fig 2A-D). Stratifying m^6^A sites by score, we observed distinct sequence patterns in highly versus lowly methylated sites. Among high-scoring m^6^A sites (score > 0.9), 93% (n = 879/945) had a cytosine immediately following the m^6^A (Fig 2C). High-scoring sites lacking a +1 cytosine were characterized by a preceding guanosine and a stretch of uracils following the m^6^A site, with a +4 uracil present in nearly all sequences (Fig 2D). This suggested that a +4 U supported an efficient m^6^A installation (Fig 2D). Indeed, low-scoring m^6^A sites with U at the +1 position exhibited a lower frequency of uracil at the +4 position (Fig S2A-B; Table S1).

**Figure 2.**
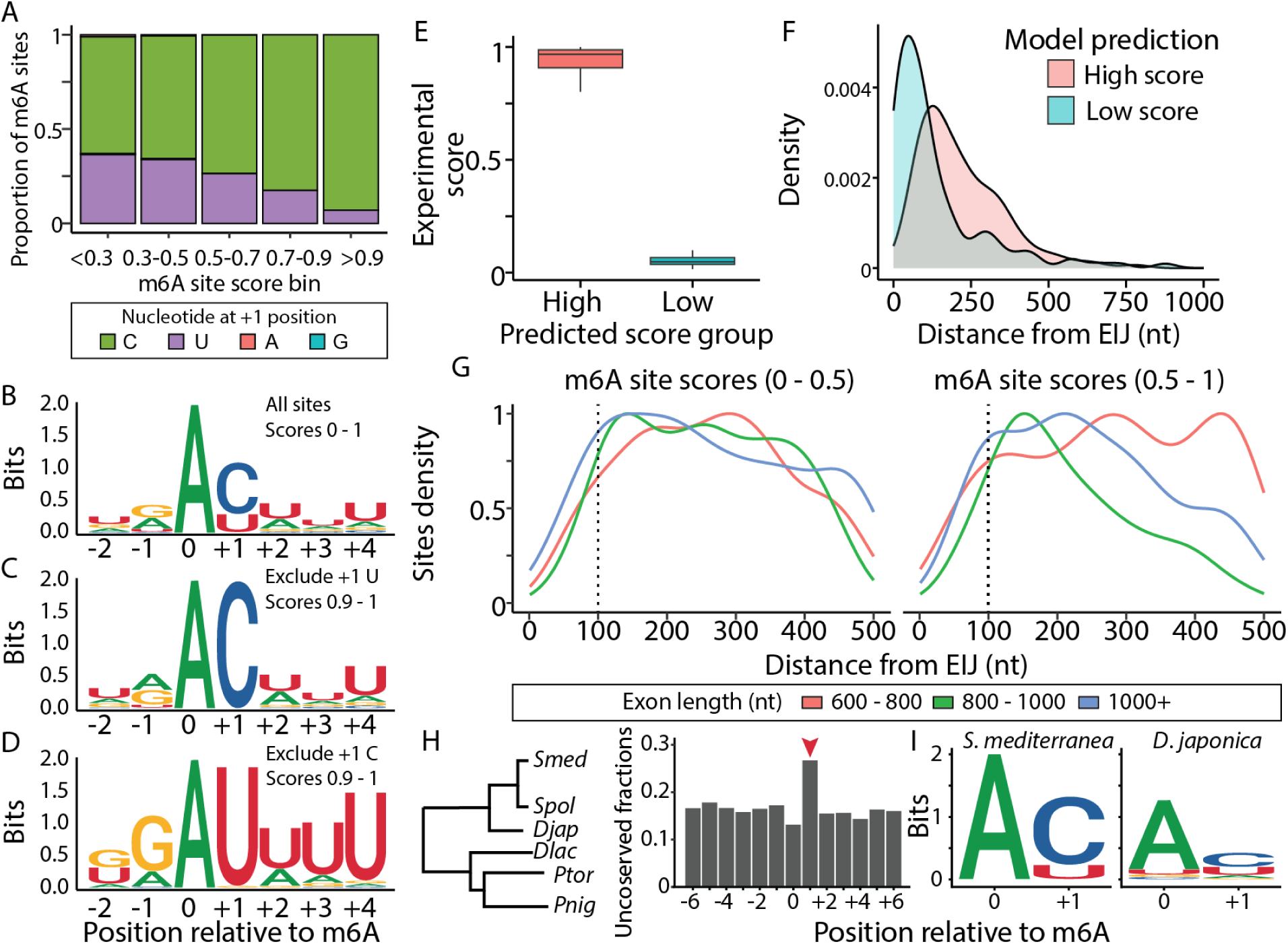
Sequence determinants of planarian m^6^A sites. (A) Examination of the nucleotide at the +1 position relative to the m^6^A site shows that regardless of the m^6^A score, the nucleotide is either C or U. Moreover, frequently modified sites are characterized by having C at the +1 position. (B-D) Consensus sequences at the m^6^A sites show that. Sites having C at the +1 position are not characterized by additional sequence characteristics (C). By contrast, strong m^6^A sites having U at the +1 position are characterized by additional sequence characteristics, with strong preference for a U at the +4 position (D). (E) Experimentally determined m^6^A scores, which have similar sequence properties, according to a gradient boost model analysis, indicating that sequence independent factors have a major role in determining the likelihood of m^6^A installation at a site. (F) The distance from the EIJ distinguished between similar sequences that differ in their experimentally determined m^6^A score. (G) Assessment of distance of m^6^A site relative to the nearest EIJ indicates that m^6^A sites are not detectable near the EIJ, regardless of the exon length, and score of the m^6^A site (high and low m^6^A site scores, left and right, respectively). Dotted line indicates 100 nt. (H) Analysis of sequence variation in regions homologous to the detected m^6^A sites in *S. mediterranea* reveals a lack of conservation at the critical position (+1) relative to the m^6^A site (red arrow). A phylogenetic tree (left) depicting the species included in this analysis is shown (left; *Smed*: *S. mediterranea*; *Spol*: *Schmidtea polychroa*; *Djap*: *Dugesia japonica*; *Dlac*: *Dendrocoelum lacteum; Ptor*: *Planaria torva*; *Pnig*: *polycelis nigra*) adapted from PlanMine^36^. The higher substitution rate at position +1 is attributed to the increased mutation rate observed in high %GC regions (see Fig S2H). (I) Pairwise sequence comparison of m^6^A sites detected in *S. mediterranea* (left) with homologous sequences in *D. japonica* (right) indicates a general lack of sequence conservation of the m^6^A and +1 sites, suggesting divergence in m^6^A site deposition between planarian species.

m^6^A sites having either a cytosine or a uracil at the +1 position showed the same preferential localization near the 3’ end of transcripts (Fig S2C), indicating that their installation in the transcript was governed by similar principles. These findings suggest simple sequence rules for m^6^A methylation: (i) cytosine (+1) following the adenine promotes high methylation potential; (ii) A uracil (+1) can only support a low methylation frequency when not part of a uracil stretch; (iii) Adenine or guanine at (+1) were largely incompatible with methylation. We examined this observation by analyzing a rare event where alleles had sequence variations in the m^6^A motif: In one allele there was a C following the m^6^A site, and the other allele had a G (Fig S2D). Both alleles were similarly expressed in the input (untreated) libraries (Fig S2E). However, GLORI libraries revealed that transcripts with the AC allele were predominantly methylated (>90%), while those with the AG allele were not (Fig S2F). This supported the observation that guanine at the +1 position following adenine was essentially incompatible with m^6^A installation.

Our observations suggested that characteristics prevalent in high-scoring m^6^A sites (e.g., cytosine at +1 and uracil at +4) were necessary but not sufficient for high methylation potential, indicating that additional regulatory mechanisms govern m^6^A installation. To explore this, we used gradient boosting regression to model sequence features associated with m^6^A sites, and their contributions to m^6^A scores (Fig S2G-H; Methods). We then extracted m^6^A sites that were predicted to have the highest m^6^A scores by the model (Fig 2E; Top 1%; Methods), and compared sites belonging to this set, which had a high m^6^A score (>0.8; n = 134), with sites having low m^6^A score (<0.1; n = 120). We found that sites having high m^6^A score prediction but that had low experimental m^6^A site scores were located near the exon intron junction (EIJ; Fig 2F; median distance = 86.5 nt). Sites having both high predicted and experimental m^6^A scores were positioned further away from the EIJ (Fig 2F; median distance = 194 nt). These results were consistent with recent findings that EIJs are strong predictors of reduced methylation frequency ^22^. A strong depletion of m^6^A sites near EIJ further corroborated this observation (Fig 2G).

We observed that m^6^A installation follows simple sequence rules (e.g., +1 C), and requiring sufficient distance from the EIJ. This suggested that m^6^A sites may be readily gained or lost during evolution by minimal sequence changes. By contrast, if a specific m^6^A site was critical for function, the underlying sequence would be expected to be highly conserved. To evaluate these alternatives, we identified potential homologous sequences of m^6^A sites in five planarian transcriptomes^36^. We compared nucleotide conservation near the m^6^A sites, and found that key positions in the m^6^A motif (e.g., +1) were not appreciably conserved to a greater extent than adjacent nucleotides (Fig 2H–I, S2H; Table S2; Methods). Interestingly, the +1 position appeared significantly less conserved compared to other nearby positions (Adjusted p-value < 2.12×10^−16^; Methods), supporting the hypothesis that m^6^A installation at a specific site was not strongly evolutionarily conserved, and hinting to potential of loss and gain. We tested whether the reduced conservation of the +1 position indicated that there is a negative selection against m^6^A sites, or alternatively a higher substitution rate of C and G nucleotides in this low % GC genome (30-35%)^38^. Indeed, mutation rate of C and G of nucleotides near the m^6^A site was significantly higher in every position tested adjacent to the m^6^A site, in pairwise sequence comparison between *S. mediterranea* and *Dugesia japonica* (Adjusted p-value < 1^−10^; Fig S2H; Methods). This analysis indicated that if selective forces act on the sequences of m^6^A sites, they are minor compared to other forces shaping genome sequence identity (e.g., bias toward certain GC content).

Our recent functional analysis of the planarian MTC and, similarly the nuclear *ythdc-1*, has revealed that it is required for the production of intestinal progenitors, and for repressing the emergence of cells expressing neural progenitor-associated genes ^4^. To investigate the cell type specificity of m^6^A methylation, we compared GLORI methylation profiles with the planarian single-cell gene expression atlas^39^. Transcripts with high-scoring m^6^A sites were detected in all major cell types (Table S1), suggesting that m^6^A likely has additional roles in cell types not previously examined^4^. This finding suggests that elucidation of m^6^A function cannot focus exclusively on analysis of the MTC, as the lethal intestine phenotype appearing following its inhibition^4^ could mask m^6^A functions in other cell types. Instead, investigating m^6^A readers, their regulation, and expression in different tissues, could offer more targeted insights into the regulatory roles of m^6^A in planarians.

### Planarians have an expanded repertoire of YTHDF proteins

We identified planarian genes that encode potential m^6^A readers by searching sequences that putatively encode sequences similar to the conserved YTH domain by protein domain analysis (Fig 3A; Methods). This analysis identified five genes encoding planarian YTHDF proteins, in contrast to *Drosophila melanogaster*, which has a single YTHDF-encoding gene, and vertebrates, which encode three^6,18^. This finding suggested that there was an expansion of the YTHDF family in planarians compared to other animals (Fig 3A-B, S3A-B). The putative planarian YTHDFs varied in length (366 - 665 amino acids; AA) compared to human YTHDFs (559 - 614 AA), indicating a significant divergence between the planarian and human sequences. We produced a phylogenetic tree of the YTH domains in planarians, humans, and additional representative species to assess their conservation (Fig 3A-B, S3A-B; Methods). The conservation of key residues associated with m^6^A recognition within the polypeptide suggested that these proteins may interact with similar substrates (Fig S3A). However, the phylogenetic analysis strongly indicated that vertebrate YTHDFs underwent independent duplication events, with no single-copy orthology observed between planarian and vertebrate YTHDFs (Fig 3B, S3B; Methods). We named the planarian YTHDF-encoding genes *ythdfA–E* to reflect their divergent evolutionary history, and which may also contribute to functional differences between planarian and human YTHDFs (Fig 3A-B; S3A-B).

**Figure 3.**
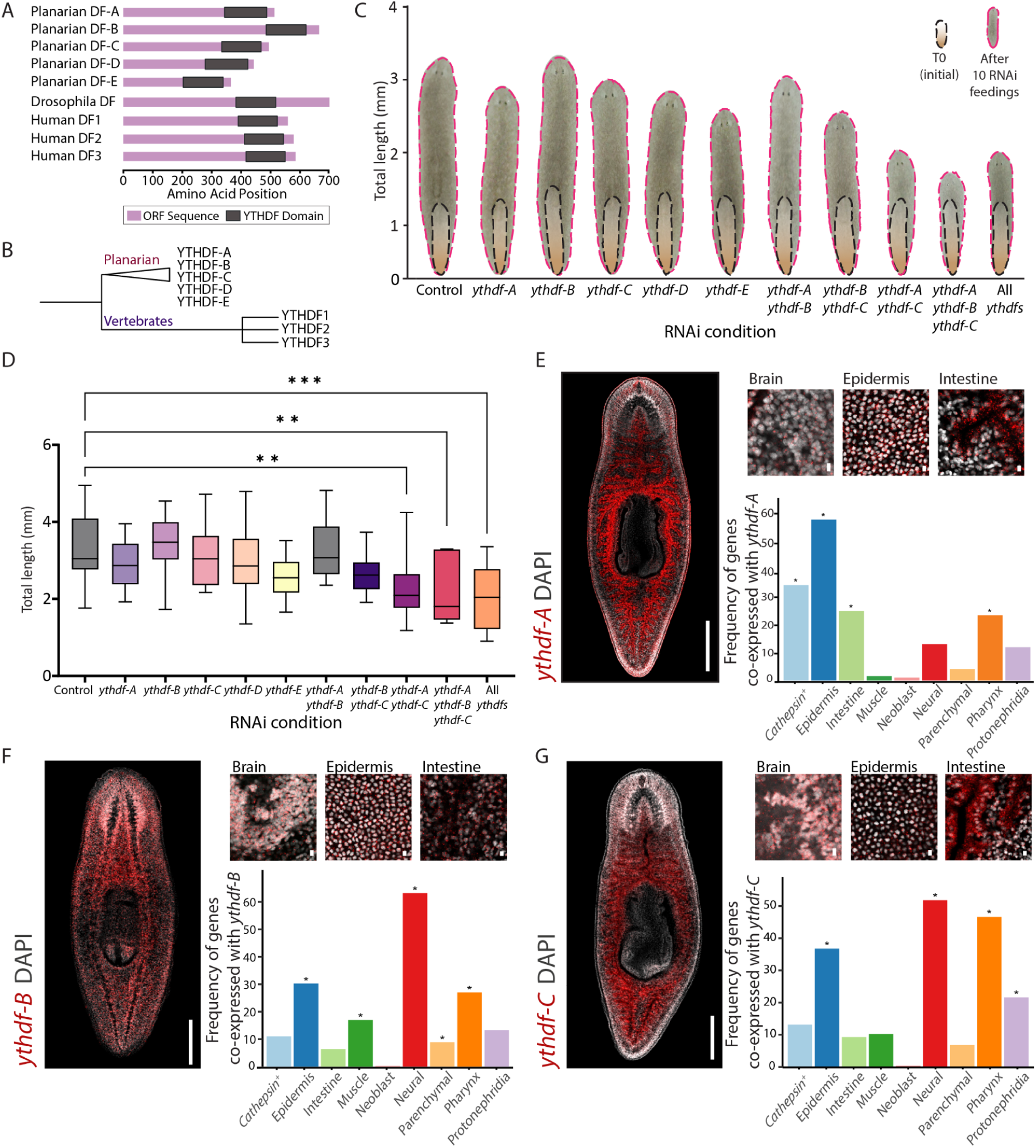
Planarians have an expanded family of YTHDF proteins. (A) A schematic representation of the YTHDF proteins across different species. The purple bars represent the protein sequence length, and the gray boxes represent the conserved YTH domain. The amino acid positions on the X-axis indicate the relative length and placement of the YTH domain within the protein sequence. While the YTH domain is highly conserved across species, the overall protein size exhibits notable variation between vertebrates and invertebrates. (B) A schematic representation of the phylogenetic analysis of YTHDF proteins (see also Fig S3B). The analysis indicated that planarian YTHDF proteins form a distinct clade from vertebrates, reflecting independent duplication events. Vertebrate YTHDF proteins cluster separately, with no evidence of orthology between planarian and vertebrate YTHDFs. (C) Inhibition of multiple *ythdf* genes results in a size reduction phenotype. Animal size was measured at the beginning of the experiment (black) and one week following the last feeding (pink). The average size of the animals is shown before and after the experiment (n>10). (D) Animal size measurements following ten RNAi feedings show a significant reduction in animal size when inhibiting at least two *ythdf* genes. Significance was calculated by using one-way ANOVA followed by Durrent’s test (**p-value < 0.01, ***p-value < 0.001). (E-G) Analysis of the expression of *ythdf-A* (D), *ythdf-B* (E), *ythdf-C* (F) by FISH and scRNAseq. Left panels show FISH analysis on the entire organism (scale = 500 µm). Right-top panels show a higher magnification at particular tissues (scale = 10 µm). Right-bottom panels show the frequency of *ythdf* gene co-expression with cell-type-specific genes, clustered by lineage origin in the planarian single cell gene expression atlas^39^. Asterisk indicates significance relative to the number of genes defining each lineage (Hypergeometric test, p-value < 0.05; Methods).

### Redundant roles of *ythdf*s in regulating planarian body size

The roles of planarian YTHDFs have not been elucidated^4^. We analyzed their functions by inhibiting their expression using RNA interference (RNAi) (Fig 3C-D; Methods). Animals were treated with double-stranded RNA (dsRNA) 10 times, and monitored for phenotypes in homeostasis and in regeneration (Fig 3C-D, S4B). We did not detect morphological or behavioral phenotypes, in either homeostasis and regeneration (Fig 3B-C, S3B; Methods). Recent studies of vertebrate YTHDF activity have demonstrated that they function redundantly and have similar biochemical targets^9,12,14^. Despite the divergent evolutionary history of planarian and vertebrate *ythdf*s, the redundancy of YTHDFs might have developed independently in planarians. We therefore tested whether planarian YTHDFs have redundant functions by co-inhibiting their expression.

We co-inhibited all five *ythdf* genes and observed a striking reduction in animal size compared to controls (Fig 3C-D; one-way ANOVA followed by Durrent’s test p = 0.003). In order to pinpoint which *ythdf* genes were driving this phenotype, we examined their expression in published scRNA-seq dataset (Fig S4A)^40^. This analysis revealed that *ythdf-A*, *ythdf-B*, and *ythdf-C* were broadly expressed, whereas *ythdf-D* and *ythdf-E* showed minimal expression (Fig S4A), suggesting a limited contribution to the phenotype (Fig 3C-D, S4B). To test this hypothesis, we co-inhibited *ythdf-A*, *ythdf-B*, and *ythdf-C* in pairs or together (Fig 3C-D, S4B), and compared the worm sizes to the inhibition of all five *ythdf* readers. The phenotype from the triple gene inhibition closely replicated the inhibition of all five genes and was stronger than any of any pair of *ythdf* genes (Fig 3C-D; S34B). qPCR confirmed over 80% inhibition in expression for each targeted gene (Fig S4C). Interestingly, despite this strong homeostatic phenotype, the animals retained their ability to regenerate (Fig S4B), in contrast to the consequence of inhibition of the MTC or of the nuclear m^6^A reader, *ythdc-1*^4^.

Suppression of the MTC causes size reduction combined with severe intestine and food ingestion defects^4^. Co-inhibition of *ythdf*s indeed resulted in size reduction, but it did not affect food uptake. Therefore, those effects might be mediated by different processes regulated by m^6^A and its mediators (e.g., *ythdc-1*^4^). To assess what cell types might be affected directly by the three *ythdf*s, which might contribute to the size reduction phenotype, we analyzed their expression by fluorescence in situ hybridization (Fig 3E-G; FISH; Methods). All three *ythdf*s were expressed in multiple tissues throughout the body showing both overlapping and distinct expression patterns(Fig 3E-G). For example, the three *ythdf*s were similarly expressed in the epidermis. Additionally, each *ythdf* had a major domain of expression in a specific organ system: *ythdf-A* in the intestine, *ythdf-B* in the brain, and *ythdf-C* in the lining of the intestine and brain (Fig 3E-G). Re-analysis of single cell RNA sequencing (scRNAseq) data from the planarian cell type-specific gene expression atlas^39^ verified that *ythdf*s are expressed across many cell types (Fig 3E-G). Moreover, the scRNAseq analysis showed that the *ythdf* genes were predominantly expressed in differentiated cells (Fig 3E-G), in contrast to the MTC components and *ythdc-1*, which are overexpressed in neoblasts^4^. This suggested that *ythdf*s function at later stages of cellular differentiation and maintenance. The co-expression of *ythdf*s was consistent with the hypothesis that the *ythdf*s may be functionally redundant.

### YTH-encoding genes are co-expressed in cells but exhibit distinct tissue enrichment

Our FISH and scRNAseq analyses showed that *ythdf-A, ythdf-B,* and *ythdf-C* were broadly expressed in multiple tissues (Fig 3D-F), suggesting that they might be co-expressed in the same cells. We performed multicolor FISH with probe pair combinations to detect *ythdf-A*, *ythdf-B*, and *ythdf-C*, and assess their co-expression (Fig 4A; Methods). First, we examined tissues that showed detectable expression of *ythdf* genes but that did not exhibit strong specificity for expression of any single *ythdf*, such as the epidermis (Fig 3D-F). We observed broad co-expression with most cells showing expression of at least two *ythdf* (Fig 4A). For example, over 60% of the epidermis cells expressed at least two *ythdf*s (Fig 4A-B; Fig S5). Examination of tissues that were particularly enriched with expression of one of the *ythdf*s (e.g., *ythdf-A* in the intestine; Fig 3E) indicated that in addition to the dominant expression of the enriched *ythdf*, other *ythdf*s were also expressed (Fig 4A). All *ythdf*s exhibited a speckle-like expression pattern, with the higher expression (i.e., tissue-enriched *ythdf*) observed as a higher density of speckles (Fig 4A).

**Figure 4.**
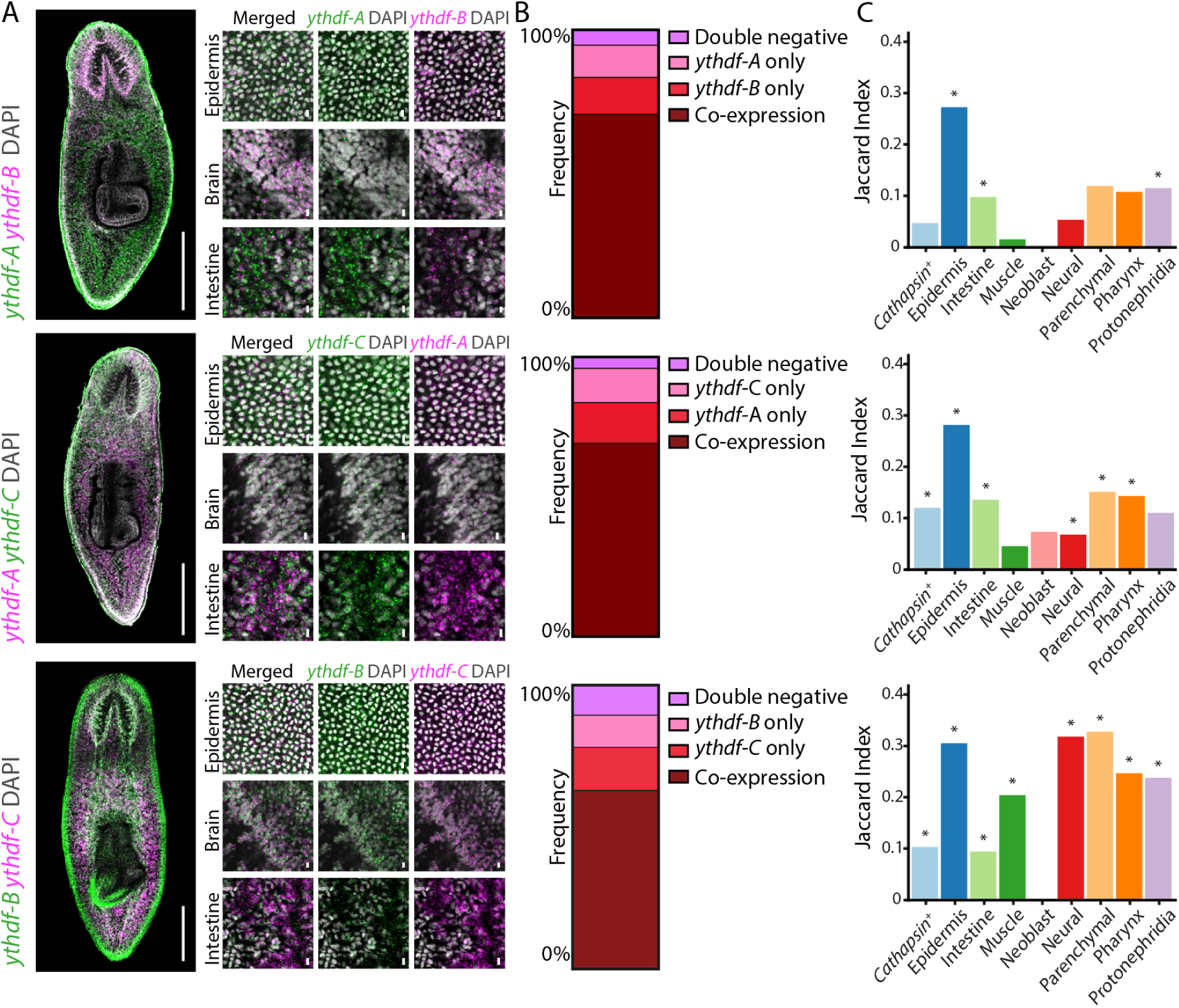
Co-expression and tissue enrichment of YTH-encoding genes. (A) Multicolor FISH of different combinations of the *ythdf* genes, showing partial co-expression. Each gene is marked with either magenta or FITC across the gene combination images (scale= 500 um) (left). Higher magnitude images of main clusters showing the differences in expression pattern of the different YTHDF proteins. (scale= 10 um) (right). (B) Proportional distribution of the expression of the *ythdf* genes in epidermal cells. Multicolor FISH analysis was performed to categorize cells into four groups: double-negative cells (light pink), cells expressing only one *ythdf* (pink and red) and cells co-expressing both *ythdf*s (dark red). Data were collected from 10 distinct epidermal regions from the top of the pharynx to the brain, normalized to the area of each region, and averaged. The percentages of each category were plotted. (C) scRNAseq analysis showing Jaccard index (i.e., overlap of co-expressed genes in tissue / union of genes; Methods) for each lineage^39^ for different pair combinations of *ythdf-A*, *ythdf-B*, and *ythdf-C*. Empirical p-value of Jaccard Index was determined using 1 M permutations. * p-value < 1×10^−4^.

To identify specific lineages expressing multiple *ythdf*s, we analyzed scRNAseq data from the planarian gene expression atlas^39^. We examined the co-expression patterns of each *ythdf* with markers associated with specific cell types (Methods). Subsequently, we quantified the similarity between the sets of genes co-expressed with different *ythdf* genes by calculating the Jaccard index for pairs of sets (Fig 4C; Methods). To determine the statistical significance of the Jaccard Index, we performed a permutation analysis with 10^6^ iterations to obtain empirical p-values of the overlap between gene sets (Methods). This allowed us to assess the overlap in co-expression profiles between *ythdf*s across cell types systematically. We found broad co-expression of *ythdf-B* and *ythdf-C* with genes representing multiple lineages, including the muscle, neural, parenchymal, pharynx, and protonephridia lineages (Fig 4C-E). The overlap between *ythdf-A* and *ythdf-B* or *ythdf-C* was primarily found for the epidermal lineage, with lower Jaccard Index detected for other cell types (Fig 4C-E). These findings suggest that YTHDF proteins might function redundantly in tissues where their expression overlap, while also exhibiting specialized roles in tissues where they were predominantly expressed. However, both FISH and scRNAseq indicated that overlap in expression of more than a single *ythdf* was prevalent in many tissues.

### Planarian YTHDFs co-regulate gene expression

The simultaneous suppression of the three *ythdf*s resulted in a significant reduction in animal size, whereas suppressing any individual *ythdf* did not produce a similar effect (Fig 3C-D). Together with the observed co-expression of *ythdf* genes, this finding suggested a potential explanation: molecular redundancy. To further investigate the molecular consequences of *ythdf* inhibition, we measured gene expression by RNA sequencing (RNAseq) following the suppression of *ythdf* genes either individually or together (Fig 5A-B; Table S5; Methods). We observed a highly significant reduction in expression of the suppressed *ythdf* gene (or genes) in each condition, with each targeted gene exhibiting a decreased expression of over 79% (Adjusted p-value < 1×10^−132^; Fig 5A-B, Fig S6A; Table S5).

**Figure 5.**
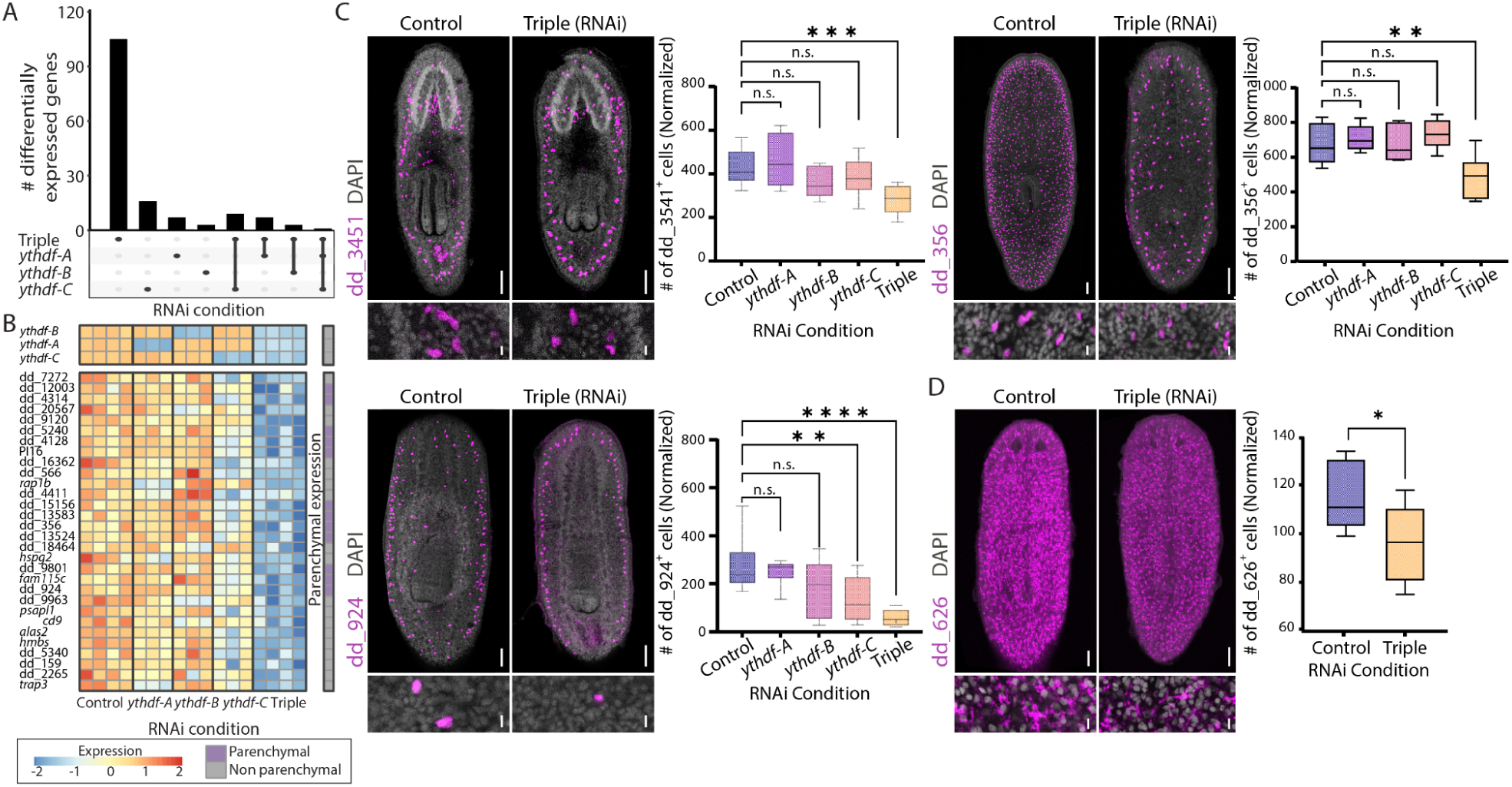
Inhibition of *ythdf* genes resulted in a decrease in parenchymal cells. (A) UpSet plot showing the number of differentially expressed genes across different *ythdf* RNAi conditions (|log_2_ (fold change)| > 0.5; Adjusted p-value < 1×10^−5^). Each bar represents a unique combination of genes shared among the specified conditions. Bars indicate the number of genes present in the intersections of selected conditions. (B) Heatmap of the top 30 downregulated genes following co-inhibition of *ythdf*s, compared to individual *ythdf* inhibition and control (FDR < 1 × 10⁻⁵). Displayed are z-scores ranging from −2 to 2. Rows represent genes, and columns represent samples. Blue and red indicate low to high gene expression, respectively. The rightmost column denotes whether the gene is highly expressed in the parenchymal lineage ^39^. (C) FISH analysis following inhibition of *ythdf* genes reveals changes in different parenchymal cell types, as described in the planarian cell type atlas ^39^. Representative FISH images are shown for animals subjected to *ythdf* co-suppression and controls. Cell counts were normalized to animal size (Methods), and compared to control animals (one-way ANOVA,*p-value < 0.05, **p-value < 0.01, ***p-value < 0.001). Scale bar = 100 μm. (D) FISH analysis detecting cathepsin⁺ cells expressing the marker dd_626 ^39^ following *ythdf* co-inhibition in RNAi treated and control animals. A comparison of normalized cell counts in the region between the pharynx and the brain (Methods) revealed a significant reduction in *cathepsin*⁺ cells in triple (RNAi) animals (p-value = 0.04, Student’s *t*-test). Scale bar = 100 μm.

Inhibiting any single *ythdf* gene did not alter the expression of other *ythdf*s or MTC encoding genes, indicating lack of compensatory gene expression within the pathway (Fig 5B, S6A; Table S5). Moreover, suppressing a single *ythdf* had a minor effect on gene expression, with the number of differentially expressed genes ranging from 6 to 26 (Fig 5A-B, Table S5; Adjusted p-value < 1×10^−5^; |log_2_ fold-change| > 0.5; Methods). Co-suppression resulted in ∼5-fold more genes significantly changing their expression, with a much stronger effect size and significance (Fig 5A; Table S5). Our results are more consistent with a model in which a differentially expressed gene is regulated, directly or indirectly, by multiple YTHDF proteins, although direct biochemical evidence is limited by the constraints of our model system (see Discussion).

We initially focused on the identity of genes that were downregulated following co-suppression of the *ythdfs* (Fig 5B). We annotated the downregulated genes using the planarian cell type atlas^39^. Analysis of the 30 most downregulated genes showed that 40% were associated with multiple parenchymal cell types^39^ (Fig 5B; n = 12/30; Adjusted p-value < 1×10^−5^; Table S5; Methods), a cell lineage giving rise to multiple secretory cell types^41^. In addition, we observed a reduction in the expression of genes active in phagocytes (n = 28/92; Table S5), suggesting an effect on either intestine or *cathepsin^+^* cell lineages, which share similar gene expression profile^39^ (Table S5).

Using FISH for detecting the expression of downregulated genes, we tested whether the gene expression reduction resulted from a decrease in the number of cells expressing the gene, or from lower expression in a comparable number of cells (Fig 5C-D, S6B-C). We found a highly significant reduction in the number of multiple parenchymal cell types following *ythdf* co-inhibition (Fig 5C, S6B), which in most cases, was not observed following inhibition of an individual *ythdf* (Fig 5C, S6B). FISH quantification of *cathepsin^+^* and intestine cells using specific markers showed a reduction in *cathepsin^+^*, but not in intestine cell numbers (Fig 5D, S6C). We note that throughout the experiment animals appeared to uptake food normally, further indicating that the intestine was not compromised. This strongly suggested that the reduction in phagocytic gene expression observed in RNAseq (Table S5) likely resulted from depletion of *cathepsin^+^* cells and not an effect on intestinal phagocytes^39,42,43^.

We assessed whether the gene expression changes that followed the *ythdf* co-inhibition were also observed after suppression of the planarian MTC^4^. We observed only a moderate correlation (R^2^ range between 0.23 - 0.4) between gene expression changes emerging following inhibition of the *ythdf*s and MTC components (Fig S6D). For example, we analyzed the published gene expression following *kiaa1429* and used the same criteria for determining the identity of the downregulated genes (adjusted p-value < 1×10^−5^; log_2_ fold-change < −0.5). Only 11 genes were similarly downregulated in *kiaa1429* (RNAi) and in the combined *ythdf* suppression (Table S5, Fig S6D). The rapid deterioration of the animal following inhibition of the MTC, which involves severe intestine damage^4^, likely masked functions mediated by these three *ythdf*s. Notably, a complementary analysis of m^6^A distribution across cell types using our GLORI data (Table S1-2) revealed no distinct enrichment in parenchymal or *cathepsin*^+^ cells. This suggested that the depletion of these cell types resulted indirectly from m^6^A regulation inactivation rather than direct targeting of m^6^A-modified transcripts.

We next examined genes that were upregulated following the inhibition of *ythdf*s (Fig 6A). Suppression of a single *ythdf* resulted in very few significant gene expression changes ranging for 1 to 12 (Table S5; adjusted p-value < 1×10^−5^; log_2_ fold-change > 0.5). In comparison, co-suppression of the *ythdf*s resulted in the upregulation of 34 genes, with 41% of the genes annotated as neural-expressed (Table S5)^39^. The inhibition of MTC components, including *kiaa1429* suppression, results in the emergence of a population of cells with a distinct neural progenitor-like gene expression profile that is almost undetectable in control animals^4^. We previously named these cells *kiaa1429* (RNAi)-specific cells^4^. Genes that are uniquely expressed in this cell population were also the most highly induced following co-suppression of the *ythdf*s (Fig 6A; Table S5). Using FISH for detection of a marker gene (dd_1837) of this *kiaa1429* (RNAi)-specific cells, we observed broad expression across the animal following co-suppression of the three *ythdf*s, beyond the normal domain of this gene’s expression around the pharynx (Fig 6B). This highly significant overabundance in dd_1837^+^ cells (Fig 6B; p = 0.0002) was especially notable in the head region, in agreement with the observation that these *kiaa1429* (RNAi)-specific cells express genes associated with neural progenitors^4^. Interestingly, these highly induced genes are found in multiple adjacent repetitive copies in the planarian genome^4^. Examination of the GLORI data mapped to this region showed that the clustered genes contain multiple m^6^A sites (Fig 6C), and that a significant, yet milder, effect was observed following the suppression of *ythdf-A* and *ythdf-C* (Fig 6A; Table S5). The presence of multiple m^6^A sites on these duplicated genes suggested that YTHDFs might recognized them, and potentially mediated their suppression in control animals.

**Figure 6.**
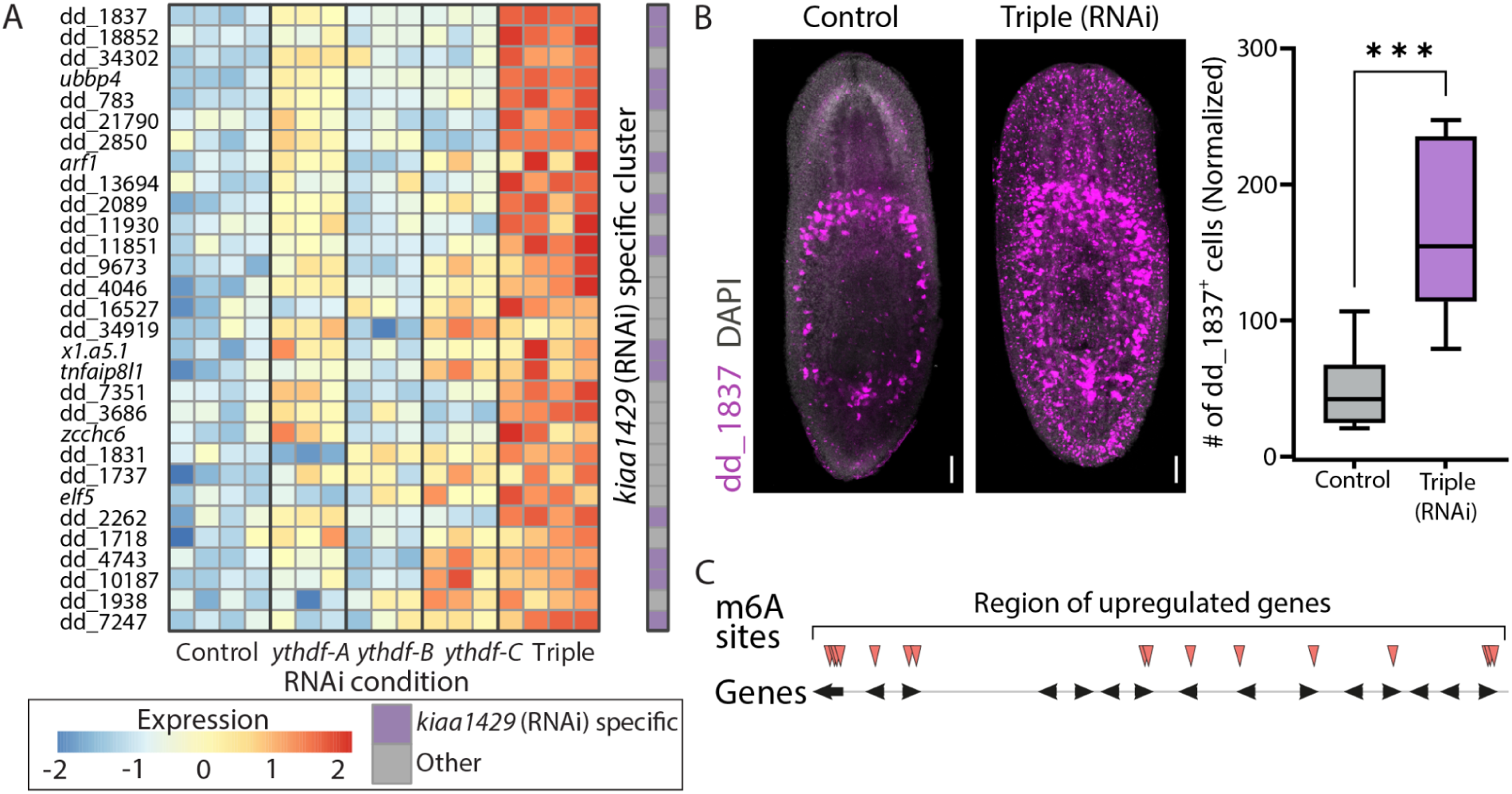
Inhibition of *ythdf* genes results in excessive production of an abnormal cell population. (A) Heatmap of the top 30 upregulated genes following co-inhibition of *ythdf*s, compared to individual *ythdf* suppression and control (FDR < 1 × 10⁻⁵). Displayed are z-scores ranging from −2 to 2. Rows represent genes, and columns represent samples. Blue and red indicate low to high gene expression, respectively. The rightmost column denotes whether the gene is highly expressed in the *kiaa1429* (RNAi)-specific cell population^4^. (B) FISH analysis detecting the *kiaa1429* (RNAi)-specific cell population marker gene dd_1837 following co-suppression of *ythdf*s and in control animals. dd_1837⁺ cells in the head region were quantified and normalized to animal size (Methods), revealing a significant increase in dd_1837⁺ cells in the RNAi animals compared to controls (Student’s *t*-test p-value = 2×10^−4^). Scale bar = 100 µm. (C) Mapping of m^6^A sites across upregulated genes in the repetitive gene cluster that characterizes *kiaa1429* (RNAi)-specific cells. Red arrowheads indicate methylation sites based on the GLORI analysis (see Table S1, S5).

### Expression of *ythdf*s is required for normal progenitor production

Previous analysis of scRNAseq data following *kiaa1429* inhibition^4^ indicated that the cell population that emerges (dd_1837^+^) has characteristics of recently produced post-mitotic progenitors, including low expression of *smedwi-1*, alongside genes associated with differentiation and lineage-specific markers^4^. To test whether the dd_1837^+^ population represents post-mitotic progenitors, we combined FISH with immunofluorescence (IF) using the SMEDWI-1 antibody, which labels neoblasts and recently divided progenitors. We observed an increased number of dd_1837^+^/ SMEDWI-1^+^ cells in the co-suppressed *ythdf*s animals compared to controls (Fig. 7A, Student’s t-test p-value < 1×10-4). To determine whether the emergence of the dd_1837^+^ progenitor population resulted from changes in the overall neoblast population, we quantified the number of SMEDWI-1^+^ cells in both control and co-suppressed *ythdf*s animals. No significant difference was found in the total number of SMEDWI-1^+^ cells between the two groups (Fig 7B). Based on these results, we hypothesize that the increase in the dd_1837^+^ progenitor-like population following co-suppression of *ythdf*s might have disrupted differentiation. Specifically, the loss of YTHDF proteins could interfere with the proper regulation of post-mitotic progenitor maturation, leading to an abnormal expansion of progenitor cells, such as the dd_1837^+^/ SMEDWI-1^+^ that fail to fully differentiate. Additionally, we propose that YTHDF proteins may play an important role in maintaining the balance between stem cell renewal and differentiation, which becomes dysregulated in their absence.

**Figure 7.**
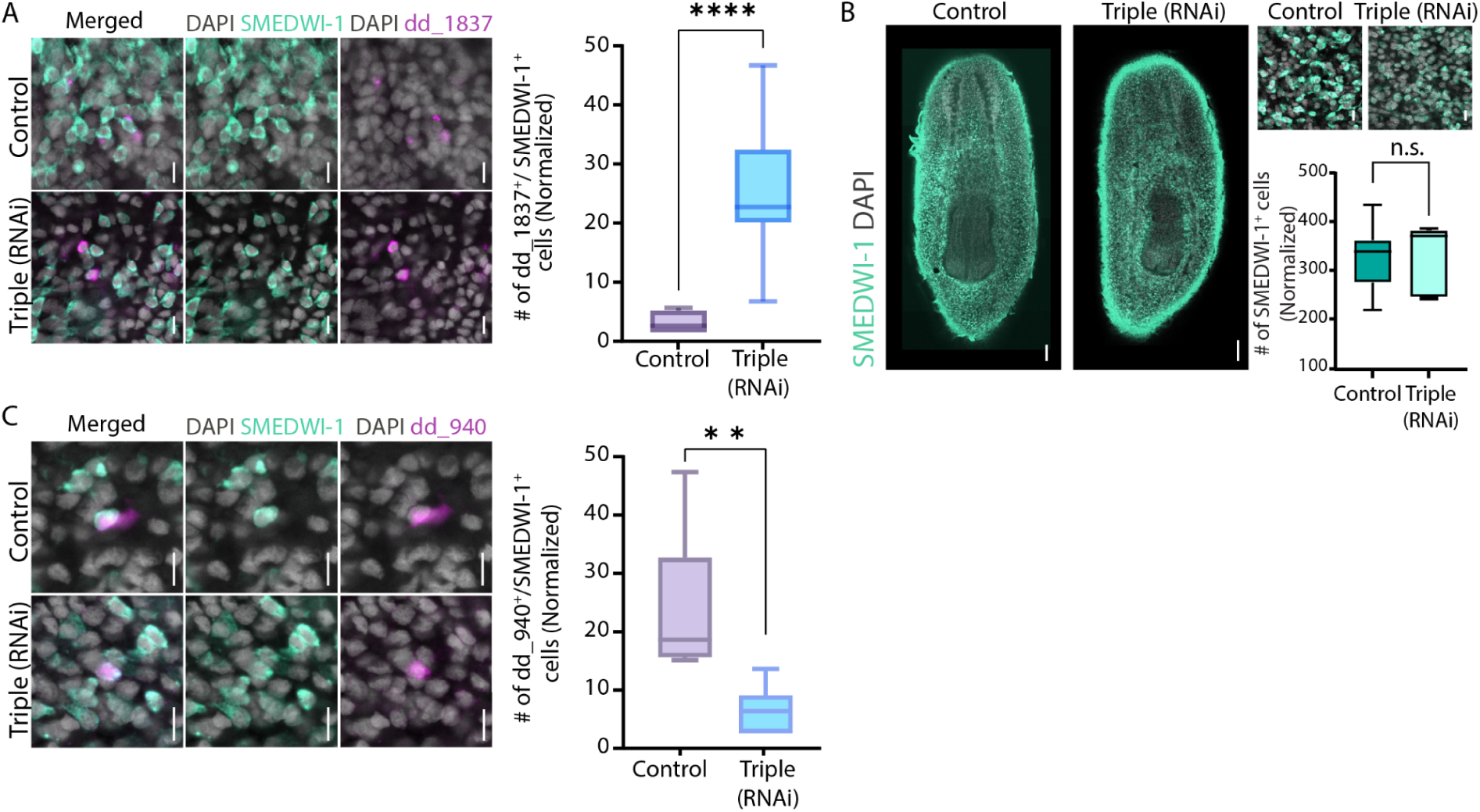
Expression of YTHDF proteins regulates the size of distinct progenitor populations. (A) Detection of dd_1837 and SMEDWI-1 by FISH and IF, respectively, is shown following co-suppression of *ythdfs* and in control animals. Cell counts of dd_1837^+^/SMEDWI-1^+^ cells in the head area were normalized to animal size and compared to control RNAi. A significant increase in dd_1837^+^/SMEDWI-1^+^ cells was observed in the triple RNAi animals (Student’s t-test, p-value < 1×10^−4^). Scale bar = 10 μm. (B) Quantification of the SMEDWI-1^+^ population size by IF is shown following co-suppression of *ythdf*s and in controls. Cell count of SMEDWI-1^+^ cells in the region between the brain and pharynx were normalized to animal size, revealing no significant difference in cell number (Student’s t-test, n.s. > 0.05). Scale bar = 100 μm. (C) Detection of the parenchymal cell marker dd_940 and SMEDWI-1 by FISH and IF, respectively, is shown following co-suppression of *ythdf*s and in control animals. Cell counts of dd_940^+^/SMEDWI-1^+^ in the region between the brain and pharynx were normalized to animal size and compared to control. A significant decrease in dd_940^+^/SMEDWI-1^+^ cells was observed in the RNAi animals (Student’s t-test, p-value = 0.0052). Scale bar = 10 μm.

To further investigate this hypothesis, we examined whether the observed decrease in parenchymal cells (Fig 5C, S6B) could be attributed to a reduction in the number of parenchymal progenitors. We combined FISH using a parenchymal marker, dd_940, combined with SMEDWI-1 IF in control and RNAi animals (Fig 7C). Quantification of double-positive cells revealed a strong reduction in parenchymal progenitors following co-suppression of RNAi condition (Fig 7C; Student’s t-test p = 0.0052), indicating that YTHDF gene expression maintained the parenchymal cell population. Together, these results show for the first time that, in planarians, effects on production of specific progenitor populations are regulated by the combined activity of YTHDFs that are likely co-expressed in the same cell, forming a hidden layer of gene expression regulation.

## Discussion

m^6^A is an essential mRNA modification in diverse biological systems^10^. In planarians, studying the MTC has revealed its essential role in producing intestine cells^4^. Analysis of a gene encoding the putative m^6^A nuclear reader, *ythdc-1*, has revealed nearly identical functions^4^. Yet, the presence of multiple other m^6^A pathway readers^4^, together with the widespread abundance of m^6^A on planarian mRNA, suggests that the pathway has additional roles, in other cell types and contexts.

Previous analyses of m^6^A in planarians, similar to many other systems, lacked single-nucleotide resolution, making it challenging to pinpoint true m^6^A sites, especially given the short installation motifs^4,44^. Our study provides the first single-nucleotide resolution map of m^6^A in planarians. Analysis of the detected m^6^A sites reveals that m^6^A deposition in planarians is governed by relatively simple sequence determinants, with strict requirements that distinguish compatible from incompatible sequences. We found that adenosines followed by cytosine at the +1 position are robust candidates for methylation. By contrast, when uracil was found at the +1 position, additional sequence elements (e.g., +4 U) were required to facilitate m^6^A installation. Certain sequences, particularly purines at the +1 position, were completely refractory to methylation. The fact that many bona fide m^6^A sites diverge from the canonical DRACH motif ^6^ might reflect an adaptation of the m^6^A pathway to the low (30%) GC content of planarian genomes^45^. Our single nucleotide resolution analysis of m^6^A indicated that methylation of adjacent sites was frequently mediated as independent events. Despite the lack of linkage in m^6^A installation, the moderate correlation between m^6^A scores of nearby sites, indicated that additional local characteristics of the transcript (e.g., general accessibility to the MTC) contributed to methylation level^22^. This model of m^6^A deposition suggested that each site was regulated as an autonomous unit. Therefore, the functional impact of m^6^A may be distributed over the transcript rather than being driven by individual sites. Moreover, the potential lack of direct association between methylation of sites, suggested that genes can gain or lose m^6^A sites, without impacting the potential functionality of remaining sites. Gain and loss of sites can occur rapidly through single-nucleotide mutations (e.g., +1 C to +1 G). This scenario supports an evolutionary model in which selective pressures for retaining m^6^A sites in a gene could act primarily at the level of transcript function rather than by strict nucleotide conservation.

In addition to installation, our work highlights the role of m^6^A in regulating cellular processes. In vertebrates, the three YTHDF paralogs appear to have highly redundant functions in regulating m^6^A-modified transcripts, predominantly facilitating mRNA turnover^9,12,14^. In planarians, there are five YTHDF homologs, which we found to have an evolutionary history distinct from their vertebrate counterparts. These planarian *ythdf*s are lowly but broadly expressed^4,39,41^, and their overlapping expression patterns hint at functional redundancy. Although techniques like CLIP-seq would be ideal for assessing in vivo binding affinities^46^, such studies are currently precluded by the lack of antibodies for planarian YTHDFs; moreover, in vitro pulldown assays are unlikely to fully recapitulate the native binding dynamics. Indeed, suppression of an individual *ythdf* does not result in a phenotype. By contrast, suppression of multiple *ythdf*s caused a size reduction, and disrupted neoblast differentiation toward multiple parenchymal and *cathepsin*^+^ cell types, dynamic processes, requiring tight regulation of transcriptional programs^43,47,48^. This observation is compatible with the possibility that planarian YTHDFs may be molecularly redundant, but importantly, because of the distinct evolutionary histories (e.g., gene duplications) of planarian and vertebrate YTHDFs, such redundancy is likely the consequence of independent processes.

Comparative insights further underscore the unique features of m^6^A regulation across invertebrates. For instance, the *D. melanogaster* genome encodes a single *ythdf* gene^11^, suggesting that cytoplasmic m^6^A recognition is mediated by a single m^6^A reader. However, the absence of single-nucleotide resolution m^6^A mapping in *Drosophila* leaves open questions regarding the potential for recognition and installation of multiple methylation sites per transcript. These differences raise the possibility that while the core mechanisms of m^6^A installation might be conserved, the evolution of reader redundancy could reflect species-specific adaptations in RNA regulation.

Our findings contribute to a broader understanding of epitranscriptomic regulation in this regenerative organism. In planarians, the straightforward sequence requirements for m^6^A installation coupled with the redundant functionality of YTHDF m^6^A readers suggest a flexible mechanism to fine-tune gene expression rapidly. This flexibility is likely crucial for processes such as stem cell differentiation and tissue regeneration, where rapid shifts in transcript abundance are required. Comparative studies will likely be instrumental in revealing how variations in m^6^A regulatory mechanisms contribute to the diverse strategies for regulation of gene expression across species.

## Methods

### Sample fixation

Animals were killed with 5% N-acetyl-cysteine in PBS for 5 min, followed by fixation in 4% formaldehyde diluted in PBSTx (PBS and 0.1% triton-x) for 20 min. Animals were then briefly washed in PBSTx, incubated in a 50:50 PBSTx:methanol solution for 10 minutes and stored in methanol at −20°C until further analysis.

### Fluorescence *in situ* hybridization (FISH)

FISH was performed as previously described^49^ with minor modifications. Briefly, fixed animals were bleached with Hydrogen peroxide and Formamide for 2 hours on a light table, then treated with proteinase K (2 μg/ml) in 1x PBSTx for 10 minutes followed by fixation in 4% formaldehyde for 10 minutes. After overnight hybridizations, samples were washed twice in pre-hyb solution, 1:1 pre-hyb-2× SSC, 2× SSC, 0.2× SSC, PBSTx. blocking was performed in 0.5% Roche Western Blocking Reagent and 5% inactivated horse serum in 1× PBSTx. Animals were incubated in an antibody overnight at 4°C (anti-DIG-POD, 1:1,500; Roche, CAT11207733910). Post-antibody washes and tyramide signal amplification were carried out as described in^49^. Finally Specimens were counterstained with DAPI overnight at 4°C (Sigma, 1 μg/ml in PBSTx).

### Fluorescence *in situ* hybridization (FISH) combined with immunofluorescence

Fixed animals were rehydrated and bleached with hydrogen peroxide and formamide for 2 hours. They were then treated with proteinase K (2 μg/ml) in 1× PBSTx for 10 minutes, followed by fixation in 4% formaldehyde for 10 minutes. After overnight hybridization, samples were sequentially washed in pre-hybridization solution, 1:1 pre-hybridization:2× SSC, 2× SSC, 0.2× SSC, and PBSTx. For immunostaining, animals were blocked in PBSTB (PBS with 0.1% Triton X-100 and 0.25% BSA) and incubated overnight at 4°C with anti-SMEDWI-1 antibody (1:1500; kindly provided by Dr. Jochen Rink). After incubation, animals were rinsed in PBSTB and washed seven times over 4 hours. Then, samples were labeled overnight with a goat anti-rabbit HRP-conjugated antibody (1:300; Abcam, CAT#ab6721). Following six washes over 3 hours in PBSTB, antibody development was performed using the tyramide signal amplification (TSA) system with FITC-tyramide (1:1500), as previously described^49^. Signal inactivation was achieved by treating specimens with 1% sodium azide for 1 hour. The FISH protocol continued with blocking in 0.5% Roche Western Blocking Reagent (CAT #11921673001) and 5% inactivated horse serum (Biological Industries CAT #04-124-1A) in 1× PBSTx. Animals were incubated overnight at 4°C with an anti-DIG-POD antibody (1:1500). Post-antibody washes and tyramide development was carried out as previously described^49^. Finally, animals were counterstained with DAPI (1 μg/ml in PBSTx, Sigma; 1:5000) overnight at 4°C.

### RNA purification

Animals were collected into 700 μl TRI Reagent (Sigma; CAT #9424) and homogenized using 0.5 mm zirconium beads in a bead-beating homogenizer (Allsheng; Bioprep-24) for two cycles of 45 seconds each and 15 seconds hold in between, at 3,500 RPM followed by incubation at room temperature for 5 minutes. Next, 140 μl of chloroform was added to each tube, followed by viguros shake for 15 seconds and incubation at room temperature for 3 min. Then, samples were centrifuged at 4°C, 12,000 *g* for 25 min for phase seperation. The upper phase was transferred into new tube and 500 μl of isopropanol was added. Tubes were inverted five times and incubated for 10 min at room temperature. RNA was precipitated by centrifugation at 12,000 g for 45 minutes at 4°C. The resulting RNA pellet was washed twice with 75% ethanol, followed by centrifugation at 7,500 g for 5 minutes at 4°C. After air-drying for 10 minutes, RNA was resuspended in 30 μl of nuclease-free water. RNA concentration was measured by Qubit (Invitrogen; Q33226) according to the manufacturer’s protocol.

### Molecular cloning

Planarian cDNA was syntisized from total RNA using RevertAid First Strand cDNA Synthesis Kit (Thermo Scientific™, CAT K1621). Target gene amplification was performed using gene-specific primers and the resulting PCR products were cloned into pGEM-t vector using the manufacturer’s protocol (Promega; CAT #A1360). Plasmids were delivered into *E. coli* TOP10 (Thermo Fisher Scientific) by the heat-shock method. Briefly, 100 μl of bacteria was mixed with 5 μl of each of the cloned vectors, incubated on ice for 30 min, and subjected to heat shock at 42°C for 45 seconds. Transformed bacteria were supplemented with 350 μl of SOC medium, and incubated at 37°C for one hour for recovery. Following recovery, bacteria were plated on agarose plates containing 1:2,000 Ampicillin, 1:200 Isopropylthio-b-D-galactoside (IPTG), and 1:625 5-bromo-4-chloro-3-indolyl-β-D-galactopyranoside (X-gal). Plates were incubated overnight at 37°C. Sequences of purified plasmids were validated by Sanger sequencing.

### Synthesis of dsRNA for RNAi experiments

Double-stranded RNA (dsRNA) was synthesized as previously described^50^. Briefly, in vitro transcription (IVT) templates were prepared by PCR amplification of cloned target sequences using primers with 5′ flanking T7 promoter sequences. dsRNA was synthesized using the TranscriptAid T7 High Yield Transcription Kit (Thermo-Fisher; CAT#K0441). Transcription reactions were incubated overnight at 37°C, followed by treatment with RNase-free DNase for 20 minutes. RNA was purified by ethanol precipitation and resuspended in 25 μl of double-distilled water (ddH₂O). The integrity of dsRNA was assessed on a 1% agarose gel, and its concentration was quantified using a Qubit 4 fluorometer (Thermo Scientific), ensuring a final concentration above 5 μg/μl. For RNAi experiments, animals were starved for at least 7 days before dsRNA feeding. Animals were fed a mixture of dsRNA and beef liver in a 1:2 ratio twice a week.

### Whole-mount in situ hybridization for detection of *ythdf* expression

Whole-mount in situ hybridization was conducted as described previously^49^ with a few modifications: In brief, worms were euthanized in 5% NAC in PBS, fixed in 4% PFA in 50% PBS containing 0.15% Triton X-100 and dehydrated in 100% MeOH. After rehydration in PBS containing 0.3% Triton X-100 (PBSTx), worms were bleached in a bleaching solution (1.2% H2O2, 5% formamide, 0.5XSSC in H2O) on a light table for ∼1 h and treated with 2 μg/mL Proteinase K (NEB) in PBSTx for 10 min followed by post-fixation in 4% PFA for 10 min. The samples were placed in Pre Hybe (50% Formamide, 5X SSC, 1X Denhardts, 100 μg/μL Heparin, 1% Tween 20, 1 mg/mL torula yeast RNA, 50 mM DTT) at 58°C for 2 h and then in Hybe (50% Formamide, 5X SSC, 1X Denhardts, 100 μg/μL Heparin, 1% Tween 20, 0.25 mg/mL torula yeast RNA, 50 mM DTT, 0.05 g/mL dextran sulfate) with the RNA probes at 58°C overnight. The next day, samples were washed with Wash Hybe (50% Formamide, 0.5% Tween 20, 5XSSC, 1 X Denhardts); 1:1 Wash Hybe: 2XSSC (0.1% tween 20); 2XSSC (0.1% tween 20); 0.2XSSC (0.1% tween 20) at 58°C for 1 h (each solution). Samples were then incubated in blocking solution (5% sterile horse serum, 0.5% Roche western blocking reagent in PBSTx) for 1 h at room temperature. Following blocking, samples were incubated with antibody (anti-DIG-POD (Roche), anti-FITC-POD(Roche)) in blocking solution (1:2000 from the antibody stocks) at 4°C overnight. Fluorophore development was done through tyramide amplification by incubating samples in TSA buffer (2M NaCl, 0.1M Boric acid in H2O, pH8.5) containing 0.006% H2O2, 20 μg/mL 4-Iodophenylboronic acid and tyramide (Rhodamine or FAM) for 30 min at room temperature. For two-colour in situs, peroxidase activity of the first antibody was quenched by incubating in 200 mM sodium azide in PBSTx for at least 1 h at room temperature. The colour reaction of the second antibody was then developed exactly like the first one. After colour development, samples were placed in Scale S4 (10% Glycerol, 15% DMSO, 40% Sorbitol, 4 M Urea, 2.5% DABCO, 0.1% Triton X-100 in H2O) at 4°C overnight and mounted on slides the next day.

### Imaging of *ythdf*s FISH

Imaging of the fixed samples was performed using an Olympus IXplore SpinSR microscope equipped with a Hamamatsu Orca Flash4.0 V3 camera. Images were acquired using OLYMPUS cellSens Dimension 3.2 software. For overview images, stitching was conducted using Huygens Professional 24.04. Three samples were imaged per condition, capturing both dorsal and ventral overviews using an Olympus UPLXAPO 10x air objective (NA = 0.4). Then, high resolution images of the head and tail regions were obtained using an Olympus UPLXAPO 20X air objective (NA = 0.8).

### Cell counting in microscopy images

Fluorescence and confocal images were acquired using a confocal microscope (Zeiss LSM800), and live images were taken using a stereomicroscope (Leica S9i). Cell counting was performed on images of animals captured from either the dorsal or ventral side, depending on the gene expression pattern. Sections for counting were imaged using a z-stack mode with 4 μm slices, spanning from the intestinal lumen to the epidermis. Cell quantification using gene expression markers was conducted on at least 10 animals per condition. Counting included the region anterior to the pharynx, with the number of cells normalized to the quantified area. Parenchymal cell counts were performed on a z-projection of the entire animal, while markers for other cell types were quantified slice by slice within a defined region covering an area indicated in the figure legend. Images were processed using Adobe Photoshop, and masks of planarian outlines were defined using the lasso and object selection tools. The background color outside the defined mask was standardized in Adobe Photoshop or Adobe Illustrator. Cell counting and analysis were performed using ImageJ.

### RNA Quantification, cDNA Synthesis, and RT-PCR Analysis

RNA concentration for each sample was measured and normalized using a Qubit 4 fluorometer (Invitrogen, Q33226). cDNA synthesis was performed on 1 μg of RNA from each sample using the RevertAid H Minus First Strand cDNA Synthesis Kit (Thermo Scientific, CAT# K1631) according to the manufacturer’s protocol, using PolydT primers. Target gene expression was measured using the QuantStudio 3 Real-Time PCR system (Applied Biosystems), with two technical replicates per sample and at least two biological replicates per condition. The relative gene expression fold-change was calculated using the ΔΔCt method, with *gapdh* expression used as the endogenous control. Forward and reverse primers for each target gene were designed, and primer efficiency was tested prior to the experiment using the standard curve method, employing five cDNA concentrations. Analysis was conducted via the Thermo Fisher Cloud Platform for data processing and visualization.

### Phylogenetic analysis planarian YTHDFs

Localization of YTH domains were identified across the planarian transcriptome (dd_v6)^36^ using InterPro analysis (PF04146; PTHR12357:SF65)^51^ on predicted protein sequences of the planarian transcriptome. The predicted protein sequences of YTH domains were extracted from YTHDC/DF genes from various species representing a diversity of animals: (*Homo sapiens, Mus musculus, Drosophila melanogaster, Danio rerio, Schmidtea mediterranea, Polycelis nigra, Dugesia japonica, Macrostomum lignano, and Nematostella vectensisi, Capitella teleta, Hydra vulgaris, Echinococcus multilocularis*). For sequences published in UniProt (Table S3), the YTH domain was extracted manually from the “Family & Domain” section. The predicted aminoacid YTH domain sequences were aligned using MAFFT^52^ with an iterative refinement method L-INS-i. Then, phylogenetic analysis was performed by maximum likelihood-based inference using PhyML^53^ with parameters: [non-parametric bootstrap analysis with 100 replicates, number of relative substitution rate categories 8, and LG as the substitution model]. The resultant phylogenetic tree was visualized with iTOL^54^.

### RNA library preparation and sequencing

RNA sequencing libraries for Triple (RNAi) and their control samples were prepared using 1 µg of total RNA. mRNA was isolated using NEBNext® Poly(A) mRNA Magnetic Isolation Module (NEB-E3370) and library preparation was performed using the NEBNext Ultra II Directional RNA Library Prep Kit for Illumina (NEB-E7760S) according to the manufacturer’s protocol. Sequencing was performed on an Illumina NextSeq 550 at the Life Sciences interdepartmental sequencing unit of Tel Aviv University. Libraries were sequenced paired-end with a read length of 2×38 bp.

### PolyA RNA sequencing analysis

Paired-end RNAseq files were mapped to the planarian transcriptome (ddV6) using bowtie2 (version 2.4.1) with default parameters^55^. Gene expression matrix was produced using the featureCounts module from subread-2.0 package with parameters [-s 2 -p]^56^. Transcriptome assembly (ddv6) was obtained PlanMine^36^ and counts of isotigs corresponding to the contig were summed (_0_1, _0_2, _0_3, etc.), to a single contig in the gene expression matrix (_0). Lowly expressed genes having a sum of less than 60 reads across samples were excluded from further analysis. DESeq2 (v1.42.1) was used to determine differentially expressed genes by pairwise gene expression analysis of each condition with control samples^57^. The differential expression analysis table was annotated using data extracted from the planarian cell-type-specific gene expression Atlas^39^.

### scRNA seq analysis

Analysis was performed on data formulated by the Reddien lab^39^ (GSE111764, Principal Clustering Digital Expression Matrix). A co-expression gene list was produced using FindMarkers, where ident.1 is defined as cells with non-zero expression level for the YTHDF gene in question, and min.pct set to 0. Each gene list was filtered for genes with p-adjusted value lower than 0.05, and average-log2 fold change larger than 0.25 to analyze significant co-expressed genes for each YTHDF. Defining lineage association for each gene was performed using single cell sequencing data available through Planarian Digiworm^39^. Defining lineage association for each gene was performed using single cell sequencing data available through Planarian Digiworm^39^. Each gene was associated with a specific lineage based on the marker genes of each main/sub cluster in the annotated dataset. This was performed using merge functions in R, creating tables containing the contig ID as well as main and sub clustering annotations (Sup). For bar chart analysis of these co-expressed genes indicating the frequency of how many genes are associated with each lineage, the number of genes associated with each cluster was divided by the number of co-expressed genes in each condition. This includes genes that are not associated with any main/sub cluster, although the unknown genes were not plotted in these graphs. Statistical significance was measured using a hypergeometric test for each bar in the bar charts. The number of genes in each lineage/cluster defines the gene population (N), the number of co-expressed genes overall is the sample size (n), and the number of genes associated with each lineage/cluster from the sample size is the number of successes (k). Each time the number of successes was tested to see whether it is greater than the random variable (x) and a p-value determines significance (0.05 threshold).

Co-expressed were assessed for common genes for each YTHDF, by finding the union of genes associated to each cluster, each time for two lists of co-expressing genes of YTHDFs (A-C). Of this union, the bar charts show frequency of overlap between YTHDFs in terms of the co-expressed genes themself, by taking the number of genes for each cluster that are in common for the two/three YTHDFs and dividing by the union. This was performed for each lineage, followed by statistical evaluation that included performing a permutation analysis. For each YTHDF co-expression list being analyzed, we determined the number of genes associated with a specific lineage. We then randomly selected, with 106 iterations, genes from the pool of all genes linked to that lineage, ensuring that the size of each random set matched the number of genes in the original list. We repeated this process for pairs or the triplet of YTHDF (A-C) co-expression lists, comparing the number of shared genes between the original lists to the overlap observed in the 106 random iterations. A p-value was then calculated to determine whether the overlap in the original lists was significantly greater than expected by chance, indicating a non-random association between the gene lists.

### GLORI treatment to RNA

For the GLORI protocol, 100-150 ng of double poly-A selected RNA per sample was used. RNA was fragmented to an approximate size of 300 nt using the Invitrogen RNA Fragmentation Reagents kit (Invitrogen). Cleanup with Dynabeads MyOne Silane was performed after each step, except those where the sample was eluted in water. DNase treatment and dephosphorylation were conducted by incubating each sample with T4 Polynucleotide Kinase (NEB), TURBO DNase, and FastAP (Thermo Scientific) for 30 minutes at 37°C in 5× FNKBuffer (1:1:2 ratio of T4 PNK buffer:FastAP buffer:H2O). Subsequently, 3′ RNA barcode adapter ligation was performed using 100 pmol (See table below) and 36 U of T4 RNA ligase (NEB) for 1.5 hours at room temperature (23°C) for each sample. After the 3′ adapter ligation, all samples were pooled, with 20% of the pool retained as a control-RNA sample. The remaining 80% of the pooled RNA underwent GLORI treatment as described by Shen et al^29^. This involved deamination of the RNA with glyoxal and sodium nitrite, followed by RNA deprotection and purification as outlined in the published protocol^29^. cDNA synthesis for both control and treated pools was carried out using the rTd reverse transcription primer (AGACGTGTGCTCTTCCG) and SuperScript III Reverse Transcriptase (SuperScript III, 18080051). After synthesis, residual rDT primer was removed by adding 3 μL of ExoSAP-IT (Affymetrix, 75001) and incubating at 37°C for 12 minutes. RNA hydrolysis was then performed in 1 M NaOH at 70°C for 12 minutes. For 5′ adapter ligation, 50 pmol of 5iLL-22 DNA adapter (/5Phos/ANNNNNNAGATCGGAAGAGCGTCGTGTAG/3ddC/) was ligated with 45 U of T4 RNA Ligase 1 (NEB, M0437M) for 4 hours at room temperature. PCR amplification of cDNA was conducted using the KAPA HiFi PCR Kit (KAPA Biosystems KK2601). The resulting amplified cDNA libraries were then purified using AMPure XP beads (Agencourt, A63881). Sequencing was performed on Illumina NovaSeq 6000.

### GLORI adapter sequences used in this study

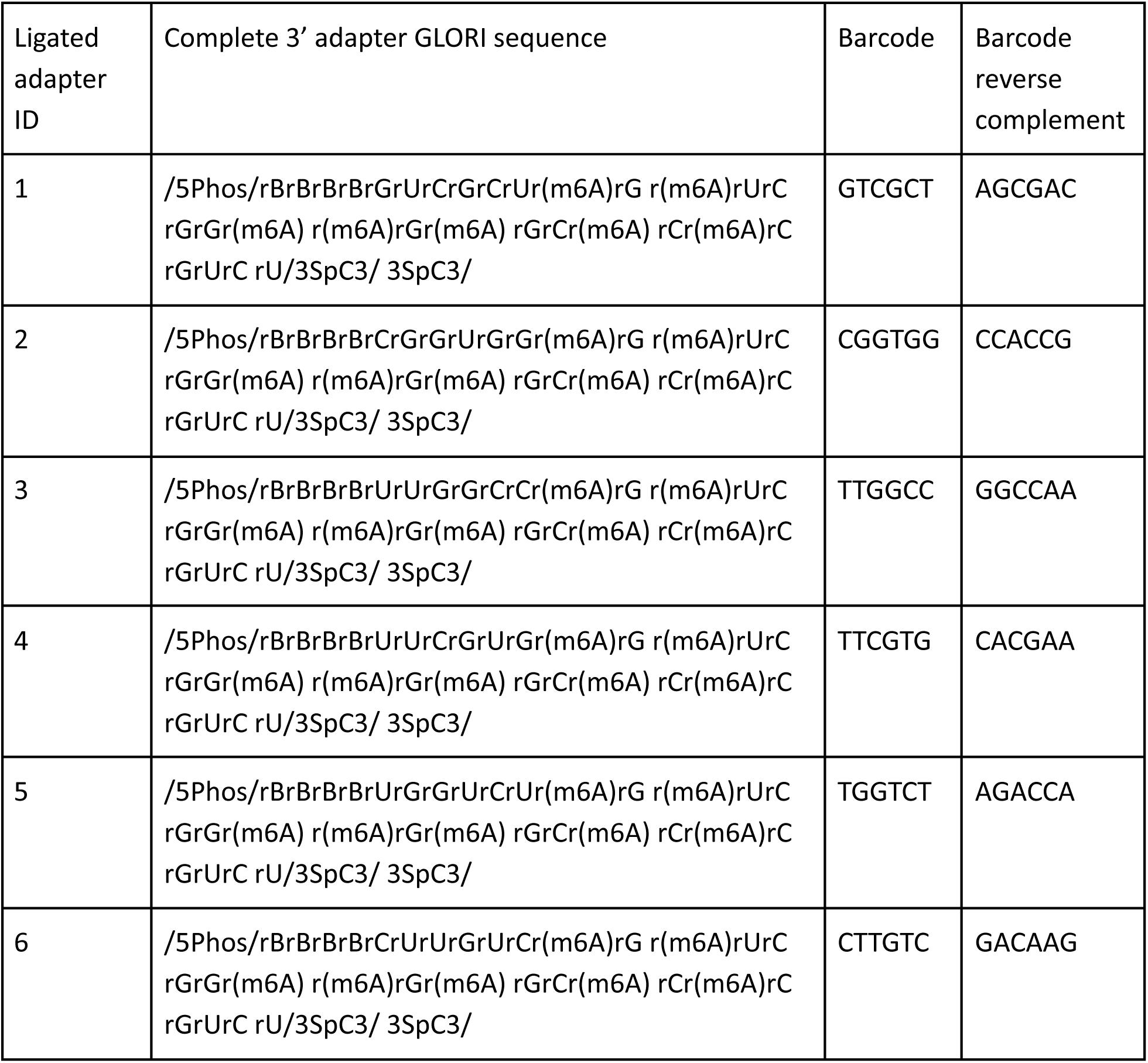

| Ligated adapter ID | Complete 3' adapter GLORI sequence | Barcode | Barcode reverse complement |
| --- | --- | --- | --- |
| 1 | /5Phos/rBrBrBrBrGrUrCrGrCrUr(m6A)rG r(m6A)rUrC rGrGr(m6A) r(m6A)rGr(m6A) rGrCr(m6A) rCr(m6A)rC rGrUrC rU/3SpC3/ 3SpC3/ | GTCGCT | AGCGAC |
| 2 | /5Phos/rBrBrBrBrCrGrGrUrGrGr(m6A)rG r(m6A)rUrC rGrGr(m6A) r(m6A)rGr(m6A) rGrCr(m6A) rCr(m6A)rC rGrUrC rU/3SpC3/ 3SpC3/ | CGGTGG | CCACCG |
| 3 | /5Phos/rBrBrBrBrUrUrGrGrCrCr(m6A)rG r(m6A)rUrC rGrGr(m6A) r(m6A)rGr(m6A) rGrCr(m6A) rCr(m6A)rC rGrUrC rU/3SpC3/ 3SpC3/ | TTGGCC | GGCCAA |
| 4 | /5Phos/rBrBrBrBrUrUrCrGrUrGr(m6A)rG r(m6A)rUrC rGrGr(m6A) r(m6A)rGr(m6A) rGrCr(m6A) rCr(m6A)rC rGrUrC rU/3SpC3/ 3SpC3/ | TTCGTG | CACGAA |
| 5 | /5Phos/rBrBrBrBrUrGrGrUrCrUr(m6A)rG r(m6A)rUrC rGrGr(m6A) r(m6A)rGr(m6A) rGrCr(m6A) rCr(m6A)rC rGrUrC rU/3SpC3/ 3SpC3/ | TGGTCT | AGACCA |
| 6 | /5Phos/rBrBrBrBrCrUrUrGrUrCr(m6A)rG r(m6A)rUrC rGrGr(m6A) r(m6A)rGr(m6A) rGrCr(m6A) rCr(m6A)rC rGrUrC rU/3SpC3/ 3SpC3/ | CTTGTC | GACAAG |

### Processing of GLORI libraries

GLORI library sequences were trimmed using cutadapt with parameters [--rrbs --paired -q 20 --stringency 1 -e 0.3 --length 15]^58^. Then, GLORI libraries were mapped to the planarian genome. *S. mediterranea* genome version schMedS1 downloaded from PlanMine^36^, and mapping was performed by reduced-representation alignment with HISAT-3N^59^ with parameters [--sp 1,0 --max-intronlen 1000] either in paired-end or single-end mode. Mapping files were converted to BAM with samtools v1.19^60^ and duplicates were marked by using PICARD tools MarkDuplicates function v2.21.4 [VALIDATION_STRINGENCY=LENIENT REMOVE_DUPLICATES=true CREATE_INDEX=true]. BAM files were processed using a custom R pipeline with the txtools package^61^. BAM files were loaded in 1 M read chunks using Rsamtools. Alignments were filtered by applying a scan flag that excluded duplicates (isDuplicate = FALSE), reads failing quality control (isNotPassingQualityControls = FALSE), and secondary alignments (isSecondaryAlignment = FALSE). Gene annotation was loaded from a BED file, and its corresponding genome sequence was used^36^. Transcriptomic reads were extracted and filtered to remove read mapping exceeding 700 nt and requiring a minimum of 10 reads per transcript. A data table was then generated using the txtools covNuc mode^61^, which recorded both coverage and nucleotide frequencies. For each adenine position, counts of A and G nucleotides were obtained and used to compute a GLORI score as the fraction of A of the sum of A and G; positions with fewer than 10 combined A+G counts were discarded. An overall adenine frequency was estimated, and Fisher’s exact tests were performed on positions with a GLORI score greater than 0.1 and a combined count of at least 10, with significance defined as p < 0.05.

### Metagene analysis of m^6^A position

A metagene plot was generated to visualize m^6^A site distribution along normalized gene lengths. Input data were filtered based on number of exons in gene, GLORI score, and replication support. If provided, coverage thresholds were applied. Gene lengths were normalized to a predefined scale (1000 nt). Single-exon genes were optionally retained or excluded. Sequence-based filtering was performed if genome sequences and motif constraints were specified by using regular expressions matching included or excluded sequences. Histogram or density plots were generated based on user preference.

### Sequence motif analysis of m^6^A sites

Sequences were extracted from the transcriptome FASTA file based on coordinates provided in the m^6^A sites table (Table S1). Filtering was performed based on strand orientation, GLORI score thresholds, and minimum replicate support. Sequences were converted from DNA to RNA. Additional filtering was applied using wildcard patterns to retain or exclude specific sequences, as labeled in figures. Sequence logos displaying information content were generated using ggseqlogo^62^.

### Co-occurrence linkage analysis between adjacent m^6^A sites

Pairs of adjacent m^6^A sites were selected based on three criteria: (1) sites were within 40 nucleotides of each other, (2) each site had a median score between 0.4 and 0.7 in control libraries, and (3) one of the sites in the pair was identified as dominant site. For each of the identified pairs, reads overlapping both sites were extracted from control BAM files using pysam library v0.22.1. Linkage between paired m^6^A sites was assessed to determine whether methylation at one site was associated with methylation at the other. A contingency table was constructed for every possible nucleotide combination across the paired sites, and a chi-squared test of independence was applied to evaluate linkage between methylation statuses. Results of the linkage analysis were visualized in R.

### Gradient boosting regression analysis of m^6^A site sequences

A predictive model was trained to correlate sequence features with m^6^A site scores. The analysis included filtering the dataset to retain sequences with a median control coverage of at least 40. Sequences around the m^6^A sites (6 nt upstream and downstream) were one-hot encoded using OneHotEncoder from scilearn v1.5.2. A gradient boosting regression model was utilized, and the dataset was divided into training and testing subsets to evaluate model performance, using mean squared error and R² score. The trained model was subsequently used to predict scores for the full dataset, and feature importance values were calculated to assess the contribution of individual features. Predicted scores were combined with metadata (e.g, distance of m^6^A site from EIJ, read depth coverage), for further analysis. Sequences having the top 1% percentile predicted m^6^A score were categorized based on experimental median control score into groups, “Coherent” (median control score > 0.8) and “Low score” (median control score < 0.1). Density plots were constructed in R to visualize the distribution of distances from the EIJ in the different categories.

### Sequencing data availability

All sequencing data produced here, including GLORI libraries and RNAseq gene expression data, was uploaded to the Sequence Read Archive (SRA)^63^ with BioProject accession PRJNA1226107.

### Analysis of m^6^A site conservation in five planarian species

Local conservation of m^6^A sites was analyzed using custom Python and R scripts. First, a tab-delimited file of m^6^A sites was parsed to extract 13-nucleotide local motifs and their associated median m^6^A scores. The motifs were converted from RNA (U) to DNA (T) and then back to RNA notation (T to U) prior to analysis. Homologous sequences were identified by matching site IDs from the m^6^A file to those in a homolog mapping CSV, and the corresponding FASTA files were selected based on a prefix index. For each m^6^A site, a sliding-window alignment was performed against the full sequence of each homolog. The alignment with the best Hamming score (i.e., the number of matching nucleotides) was retained. Alignments were discarded if the best score was ambiguous or if the Hamming distance exceeded a predefined threshold. Ambiguous mapping or matches with Hamming distance exceeding 3 were removed. Sequence logos were then generated using the ggseqlogo package^62^. For statistical evaluation of positional conservation, a two-sample proportion test was performed using the R function prop.test. For each position in the motif, the number of conserved nucleotides (as indicated by the match vector) was summed across all retained alignments, and the proportion of matches at that position was compared to the combined proportion of matches at all other positions. P-values were calculated and subsequently adjusted using a Bonferroni correction to account for multiple testing.

### Primer sequences of genes cloned in this study

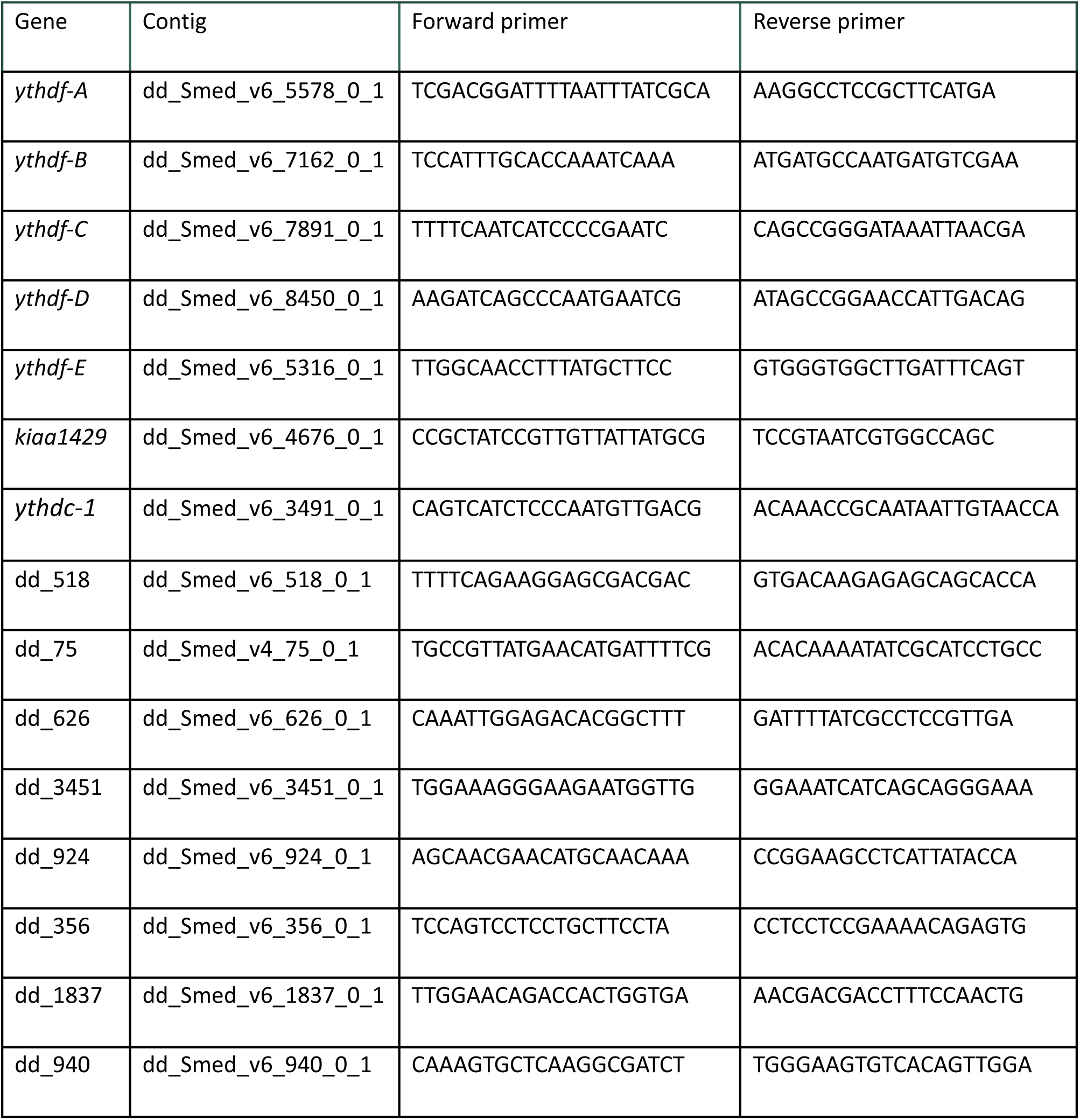

### qPCR primer table

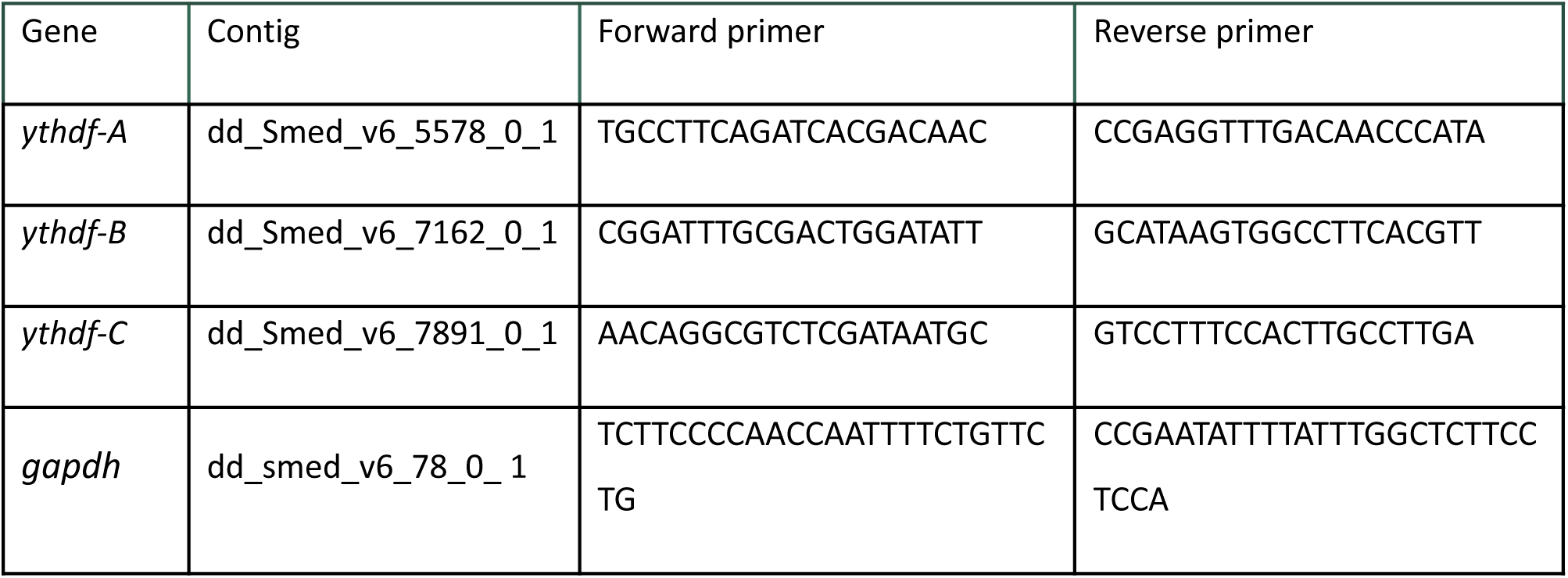

## Supporting information

Table S1. Planarian m6A site mapping across the genome.

Table S2. Planarian m6A site mapping across genes.

Table S3. Sources of protein sequences used for YTH domain phylogenetic analysis.

Table S4. Genes co-expressed with planarian ythdfs using scRNAseq analysis.

Table S5. Differential gene expression analysis following ythdfs inhibition.

Table S6. Index of contig ids corresponding to labels in figures.

## Supplementary figures

**Figure S1.**
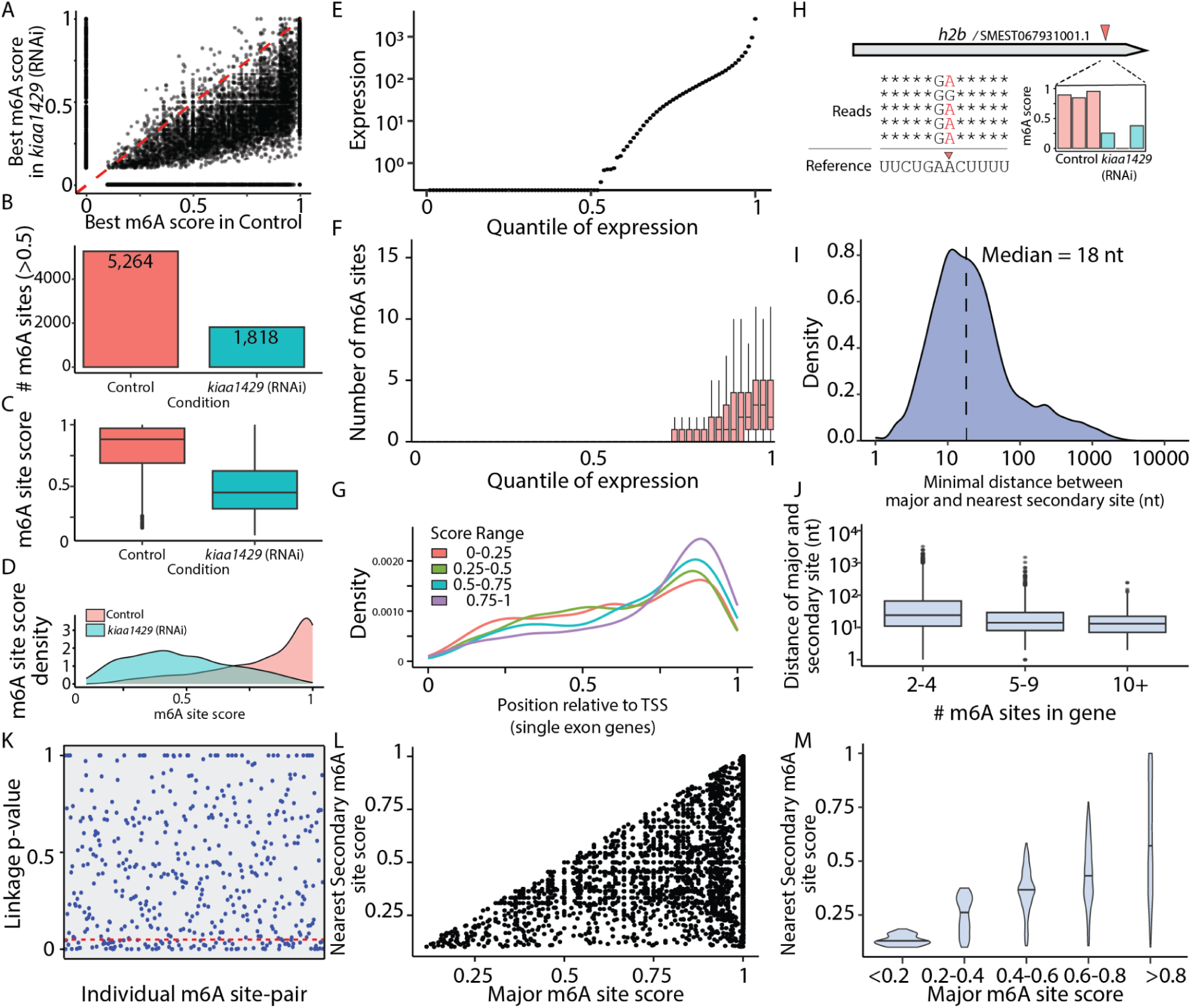
Properties of planarian m^6^A sites. (A) The scatter plot displays m^6^A site scores (black dots) in control samples, where m^6^A levels are normal, and in kiaa1429 (RNAi) samples, where m^6^A levels are reduced. The red dashed line represents a slope of one. For each site, the score shown is the highest observed across any of the three replicates for each condition. (B) Number of m^6^A sites with median score greater than the 0.5 in control and *kiaa1429* (RNAi) libraries demonstrates the reduction in overall methylation in *kiaa1429* (RNAi). (C-D) Shown is the distribution of median scores of m^6^A sites in control and *kiaa1429* (RNAi) libraries as boxplot (C) and density (D) plots, for sites having coverage of at least 10 sequencing reads (n = 2,284) in each of the three replicates used for the two conditions in this analysis. (E) Shown is the median normalized expression^57^ of each gene in input libraries as a function of the quantile of expression. For example, the median expression of ∼50% of the genes is 0. (F) The median number of m^6^A sites is shown (median score in control libraries > 0.1) as a function of quantile of gene expression, which was calculated based on the expression level of the gene in the input GLORI (i.e., untreated) libraries. (G) Metagene analysis of m^6^A positions (n = 2,609) of single gene exons shows a strong enrichment towards the 3’ end, indicating that 3’ end preference is not unlikely to be governed only by splicing specific factors. (H) Canonical histone transcripts are also targets for methylation. Shown is a canonical *h2b* (dd_2520) transcript with an m^6^A site (red arrow) detected near the 3’ end. Read sequences that cover the m^6^A site are shown (bottom left) with the methylated A highlighted in red. The m^6^A score of the site in different replicates indicates that the m^6^A installation is dependent on the MTC. (I) Shown is the minimal distance between the site having the highest m^6^A score (i.e. major) with a secondary site (i.e., m^6^A site having a lower score than the major site), in genes that have multiple m^6^A sites. (J) Shown is a comparison of distance between the major and secondary site for genes with multiple m^6^A sites. (K) The p-value of linkage of m^6^A installation is shown for m^6^A site-pairs (Methods). For each site pair (blue dot) a p-value was calculated (y-axis). (L-M) Correlation of m^6^A score between the major m^6^A site and the nearest secondary site is shown.

**Figure S2.**
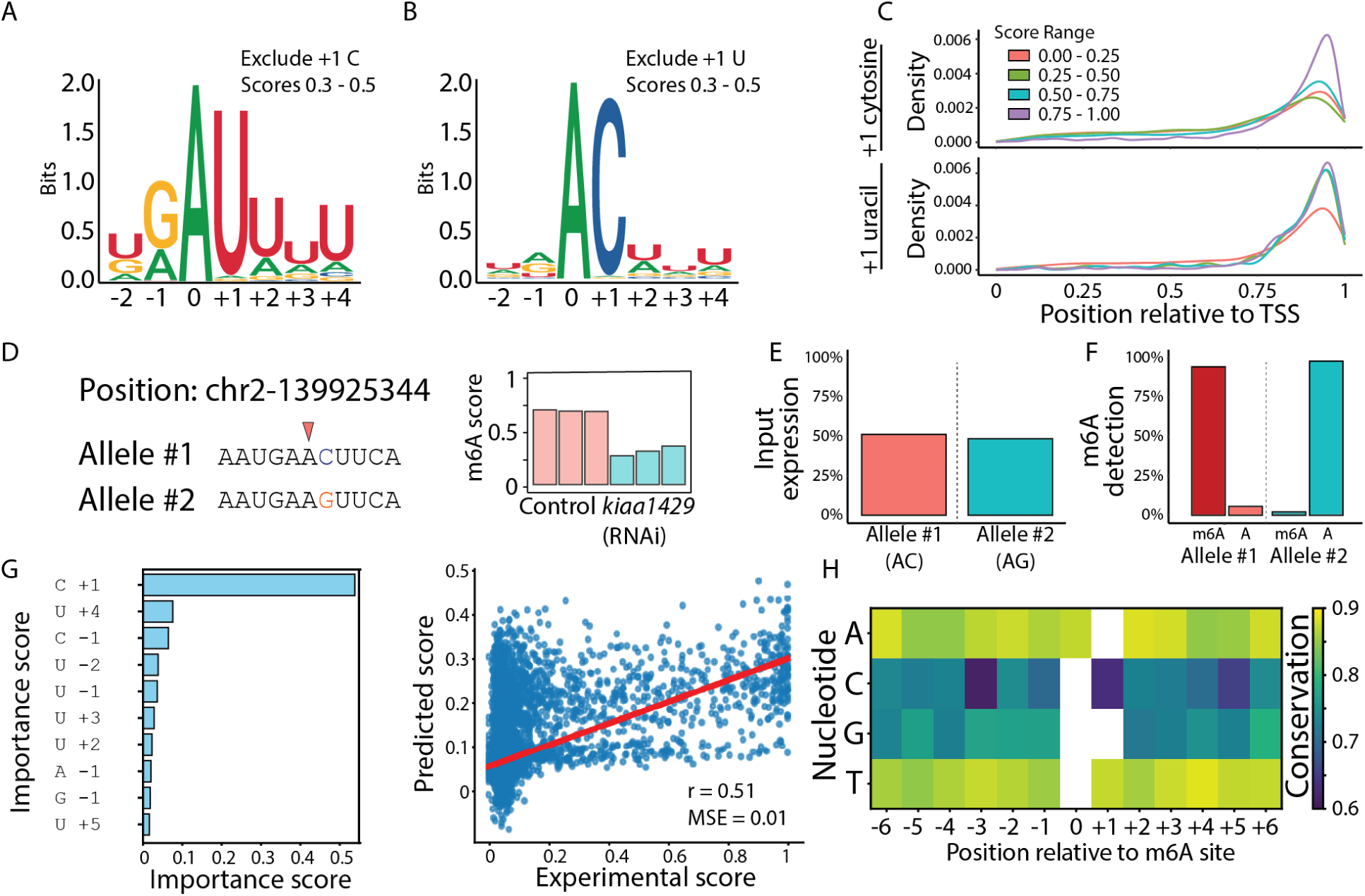
Sequence characteristics of m^6^A sites. (A) Shown is a sequence motif of m^6^A sites having low scores (0.3 - 0.5) and lacking a +1 C. The presence of a stretch of Us is observed as well as preference for a −1 G. (B) Shown is a sequence motif of m^6^A sites having low scores (0.3 - 0.5) and lacking a +1 U. There are no strong sequence preference for particular nucleotides other than a +1 C. (C) Metagene analysis of sites having either a +1 C (top) or +1 U (bottom) show a similar distribution across the gene, indicating that the sites are installed similarly, despite the incompatibility of the +1 U with the known m^6^A installation consensus DRACH. (D-F) A rare example of an m^6^A site located at a sequence position that differs between alleles is shown. (D) The site (left, red arrow) is methylated in the allele with the AC sequence, whereas the other allele, which contains a +1 G, is refractory to m^6^A installation and remains unmethylated. The m^6^A site score (right) indicates methylation in the allele containing a +1 C. (E) The expression levels of both alleles were nearly identical, as determined by counting reads covering the sequence difference and distinguishing between them based on the nucleotide identity at the +1 position relative to the m^6^A site. (F) The fraction of methylated and unmethylated A is shown for both alleles, demonstrating that a +1 G is incompatible with m^6^A deposition. (G) The importance scores of sequence features near the m^6^A site are shown. The strongest determinant of m^6^A installation is the presence of a +1 C, while a +4 U is also associated with high methylation potential (left). A model incorporating only local sequence features performed better at identifying sequences that were poor methylation targets than at predicting methylation-compatible sequences (right), suggesting that factors beyond local sequence identity influence methylation levels. (H) Sequence conservation was calculated for individual m^6^A sites between *Schmidtea mediterranea* and *Dugesia japonica*. The comparison revealed similar conservation patterns across sites when considering the nucleotide identity near the m^6^A site. Overall, C and G were less conserved than A and T.

**Figure S3.**
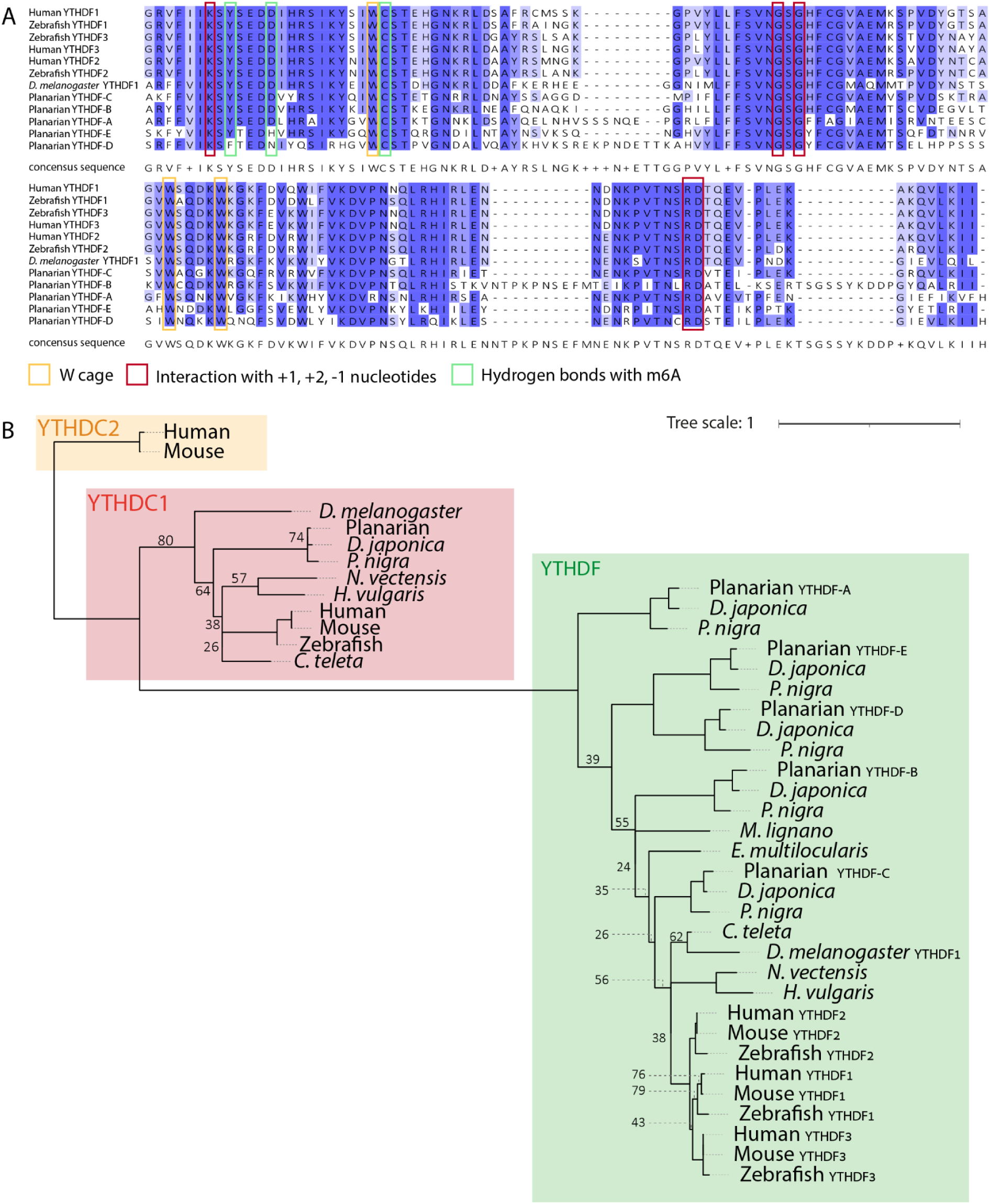
phylogenetic analysis of planarian YTHDF proteins. (A) Multiple sequence alignment of the conserved YTH-domain from YTHDF proteins in planarians, zebrafish, and *Drosophila*. Darker colors represent higher levels of amino acid conservation. (B) Phylogenetic tree illustrating the evolutionary relationships among various YTH-domain proteins across different species (Table S3). Numbers indicate bootstrap values (1-100); bootstrap values > 85 were omitted for visual clarity. Planarian: *Schmidtea mediterranea*; *Spol*: *Schmidtea polychroa*; *D. japonica*: *Dugesia japonica*; *Pnig*: *polycelis nigra; C. teleta*: *Capitella teleta*; *M. lignano*: *Macrostomum lignano*; *Echinococcus multilocularis*; *H. vulgaris*: *Hydra vulgaris*; *N. vectensis*: *Nematostella vectensis*; *D. melanogaster*: *Drosophila melanogaster*.

**Figure S4.**
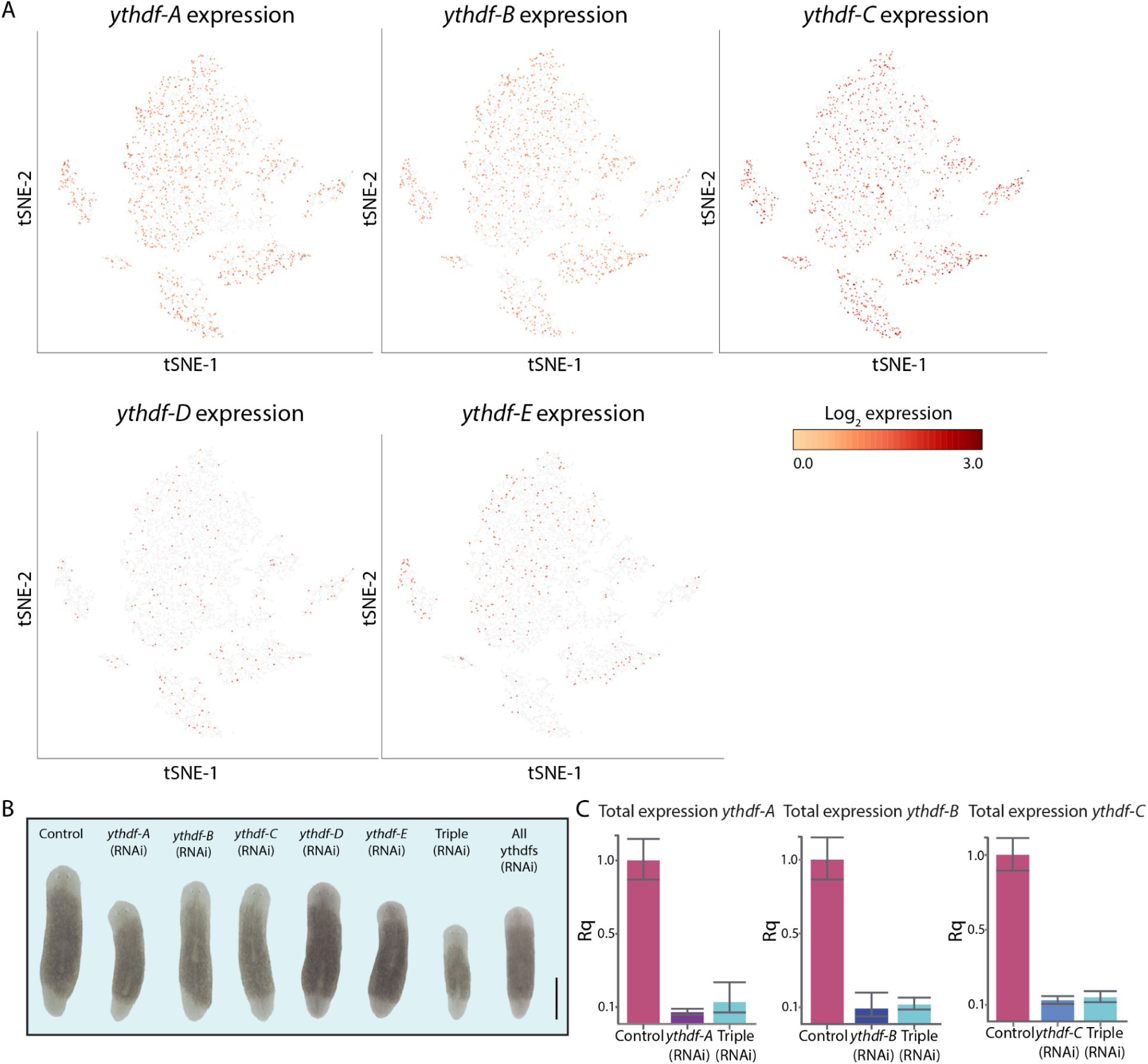
*ythdf* expression was not required for planarian regeneration. (A) *ythdf*-encoding genes exhibit broad expression across planarian cells based on scRNA-seq data. Shown here is the expression of the five *ythdf* genes, extracted from a published dataset of G0 planarian cells^40^. (B) *ythdf* (RNAi) animals successfully regenerated heads and tails following amputation (15/15). Scale bar = 1 mm. (C) qPCR analysis showing significant downregulation of *ythdf-A*, *ythdf-B*, and *ythdf-C* gene expression following RNAi in both single and triple inhibition conditions. Error bars indicate the 95% confidence interval. Data include two technical replicates and three biological replicates. p-value for each condition compared to its control was as high as 0.0078.

**Figure S5.**
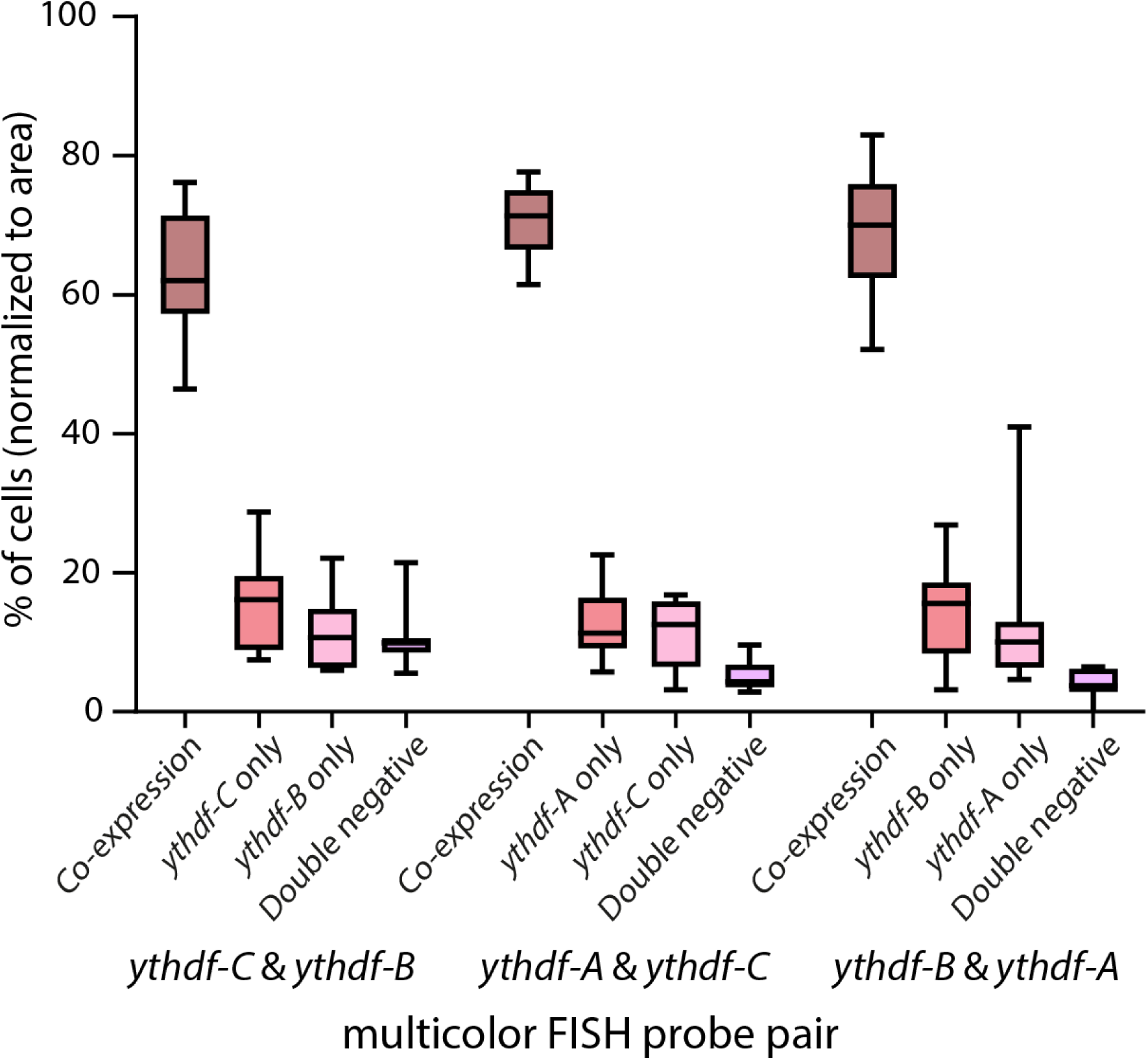
Quantification of *ythdf* expression in the planarian epidermis. A box plot represents the percentage of epidermal cells expressing *ythdf* genes. Multicolor fluorescence in situ hybridization (FISH) analysis was used to categorize cells into four groups: double-negative cells, single-positive cells (expressing only one *ythdf*), and double-positive cells (co-expressing two *ythdf* genes). Data were collected from 10 distinct epidermal regions from the top of the pharynx to the brain and normalized to the area of each region (Methods).

**Figure S6.**
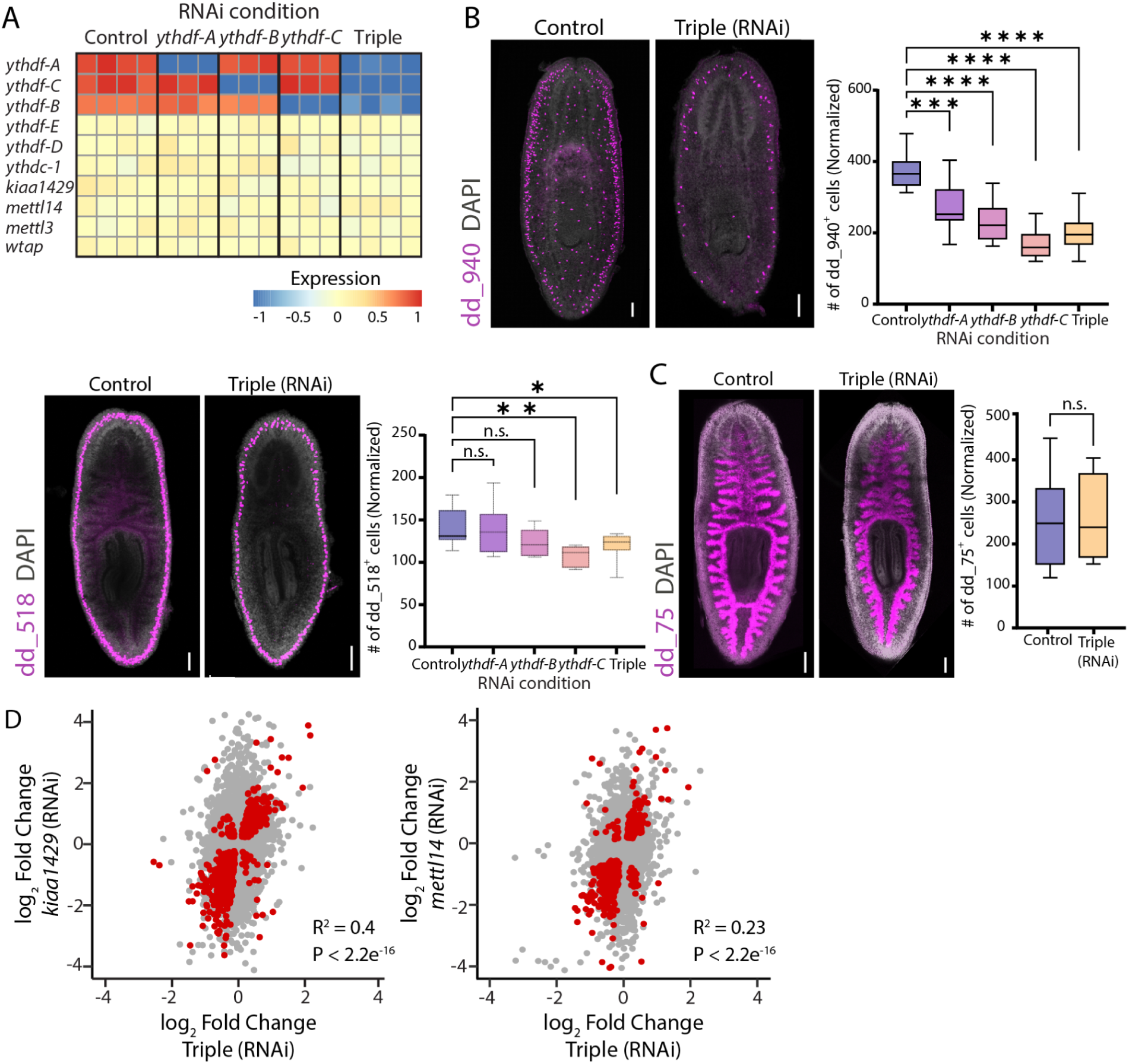
Inhibition of *ythdf* genes resulted in a reduction in parenchymal cells. (A) Heatmap of YTH-domain protein encoding genes and MTC encoding gene expression following inhibition of *ythdf* genes. Inhibition of a *ythdf* genes did not affect the expression of other components of the m^6^A pathway. Displayed are z-scores ranging from −1 to 1 (FDR < 1 × 10⁻⁵). (B) FISH analysis showing changes in different parenchymal cell types^39^ following inhibition of *ythdf* genes using the markers dd_940 (top right) and dd_518 (bottom left). Cell counts were normalized to animal size (methods) in *ythdf-A* (RNAi), *ythdf-B* (RNAi), *ythdf-C* (RNAi), and triple (RNAi) animals and compared to control animals. Statistical significance was assessed using one-way ANOVA (*p-value < 0.05, **p-value < 0.01, ***p-value < 0.001, ****p-value < 0.0001). Scale bar = 100 μm. (C) FISH analysis detecting intestinal cells expressing dd_75, in triple (RNAi) and control animals. Cell count of dd_75^+^ cells in the intestinal region spanning the pharynx was normalized to animal size, revealing no reduction in intestine cell number following *ythdf*s inhibition (Student’s t-test, p > 0.05). Scale bar = 100 μm. (D) Correlation of gene expression changes between *kiaa1429* (RNAi) and triple (RNAi) compared to their controls (left) *mettl14* (RNAi) and Triple (RNAi) compared to their controls (right). Each colored dot represents a gene, red and black represents significant (FDR < 0.05 for both conditions) and nonsignificant change in gene expression compared to controls, respectively.

Table S1. Planarian m^6^A site mapping across the genome.

Table S2. Planarian m^6^A site mapping across genes.

Table S3. Sources of protein sequences used for YTH domain phylogenetic analysis.

Table S4. Genes co-expressed with planarian *ythdf*s using scRNAseq analysis.

Table S5. Differential gene expression analysis following *ythdf*s inhibition

Table S6. Index of contig ids corresponding to labels in figures.

## Acknowledgements

We thank Hila Kobo for assistance with Illumina sequencing at the Tel Aviv University core facilities. We thank Kristina Karin Mirkes for support in confocal microscopy imaging. We thank Prof. Joel Hirsch and Dr. Aldema Sas-Chen for critical input on the manuscript. We thank the Wurtzel lab for critical input. O.W. is supported by the Israel Science Foundation (grant 2039/18) and the European Research Council (no. 853640). O.W. is a Zuckerman Faculty Scholar. A.P. is supported by the EIPOD-LinC Postdoc Fellowship from the European Molecular Biology Laboratory. H.T.K.V is supported by the European Molecular Biology Laboratory.

